# New motifs in Ndc1 mediating interaction with the Nup84 complex and nuclear membranes

**DOI:** 10.1101/2022.07.31.501833

**Authors:** Ingo Amm, Marion Weberruss, Andrea Hellwig, Johannes Schwarz, Marianna Tatarek-Nossol, Christian Lüchtenborg, Martina Kallas, Britta Brügger, Ed Hurt, Wolfram Antonin

## Abstract

The nuclear pore complex (NPC) embedded in the double nuclear membrane is built from ~30 different nucleoporins (Nups) in multiple copies, of which a few are integral nuclear membrane proteins. One of these transmembrane Nups is Ndc1, which is thought to play a role in interphase NPC assembly at the fused inner and outer nuclear membrane. In this study, we discovered a direct interaction of Ndc1’s transmembrane domain with Nup120 and Nup133, members of the Y-complex that coats the nuclear pore membrane. In addition, we identified a so far unrecognized amphipathic helix (AH) in the C-terminal domain of Ndc1, which can bind to high curvature liposomes. When overexpressed in yeast this amphipathic motif is toxic and dramatically alters the intracellular membrane organization. Further genetic investigations revealed that Ndc1-AH functionally interacts with related motifs in the C-terminus of Nups Nup53 and Nup59, known to serve in nuclear pore membrane binding and link between NPC modules. This relationship could explain why the essential function of Ndc1 can be suppressed by deleting the amphipathic helix from Nup53. Our data indicate that nuclear membrane biogenesis dependent on a balanced ratio between amphipathic motifs in diverse nucleoporins is essential for interphase NPC biogenesis.

## INTRODUCTION

Nuclear pore complexes (NPCs) are huge proteinaceous assemblies embedded in the double-layered nuclear envelope (NE). Each NPC is composed of multiples of 8 copies (8-64) of about 30 different nuclear pore proteins called nucleoporins (Nups) resulting in a total amount of about 500 up to 1000 polypeptide chains per NPC in higher metazoan. The majority of the nucleoporins are organized in biochemically and structurally defined subcomplexes: the inner nuclear pore ring connecting the NPC to the nuclear membrane, the attached central channel complex, the nuclear and cytoplasmic rings with the attached peripheral elements, the nuclear basket and the cytoplasmic filaments (Hampoelz et al., 2019; Hurt and Beck, 2015; Lin and Hoelz, 2019; Schwartz, 2016). The main function of NPCs is the selective macromolecular transport across the NE. A group of nucleoporins forming the central diffusion barrier contain natively unfolded, phenylalanine-glycine (FG)-rich sequences, blocking passive diffusion of molecules larger than ~5 nm in diameter (corresponding to a molecular mass of 30 – 40 kDa), (Frey et al., 2006; Meinema et al., 2013; Mohr et al., 2009; Popken et al., 2015; Terry and Wente, 2009). Transient interactions of the FG-Nups with distinct nuclear transport receptors (NTRs) allow active transport of large molecules and define the directionality and specificity of transport across the NE (Paci et al., 2021).

The molecular mass of the NPC up to ~120 kDa in mammals and its complex architecture raise the question of how NPCs are assembled and stabilized in the NE throughout cell cycle. Metazoan NPC biogenesis occurs in two different cell-cycle stages. The mitotic NPC assembly mode is characteristic for organisms undergoing open mitosis (Kutay et al., 2021). At the beginning of mitosis, the NE and NPCs disassemble and the NE including the nuclear membrane proteins are absorbed by the mitotic ER membrane network. In anaphase NPC reformation is initiated by the chromatin binding of ELYS/MEL-28, which recruits the Nup107-Nup160 complex to the assembling NPC. NPC assembly continues most likely into preformed openings of the ER membranes contacting the chromatin (Otsuka et al., 2018). Whereas the mitotic assembly mode in metazoan cells establishes a transport competent nucleus in daughter cells within minutes, NPC biogenesis occurring during interphase, is much slower probably due to the requirement of NPC insertion into the intact double membrane layer of the NE (Dultz and Ellenberg, 2010). In organisms undergoing closed mitosis like *Saccharomyces cerevisiae* the NE stays intact through the whole cell cycle. Thus, NPCs have to be inserted exclusively into the intact double membrane layer (Doucet and Hetzer, 2010; Otsuka and Ellenberg, 2018; Weberruss and Antonin, 2016). The major challenge is arguably bringing both the inner and outer nuclear membrane (INM and ONM) in close proximity for subsequent merger, which is an essential and rate-limiting step of interphase NPC biogenesis. In principle, membrane curvature has to be generated from both sides of the NE by applying forces to flat areas of the NE (D’Angelo et al., 2006; Doucet and Hetzer, 2010). Highly curved membranes represent high energy states characterized by unfavourable membrane lipid packing (Kozlov et al., 2010; Martens and McMahon, 2008). Curvature can be induced either by (1) NE proteins with large luminal domains bridging between INM and ONM similar to viral fusion proteins or SNAREs (soluble N-ethylmaleimide-sensitive factor attachment protein receptors), (2) insertion of amphipathic helices into the lipid bilayer, (3) formation of membrane-connected protein scaffolds or (4) generation of an asymmetric lipid composition of the two leaflets in the lipid bilayer (Antonin et al., 2008; Rothballer and Kutay, 2013). Membrane nucleoporins are good candidates of being involved in the assembly of the NPC into the intact NE. Yeast expresses four transmembrane nucleoporins (Ndc1, Pom34, Pom152 and Pom33) of which Ndc1, Pom34 and Pom152 form a stable transmembrane nuclear pore subcomplex and anchor the inner ring of the NPC in the equatorial plane of the nuclear pore membrane (Chadrin et al., 2010; Chial et al., 1998; Lau et al., 2004; Madrid et al., 2006; Onischenko et al., 2009; Rout et al., 2000; Upla et al., 2017; Wozniak et al., 1994). Ndc1 is the only known conserved membrane nucleoporin revealing sequence homology between lower (such as *Chaetomium thermophilum*) and higher eukaryotes. The C-terminal region of Ndc1 is predicted to be exposed to the cytoplasm/nucleoplasm (Lau et al., 2006; Mans et al., 2004; Mansfeld et al., 2006; Neumann et al., 2010; Stavru et al., 2006), (Fig. S2A-B). In yeasts, Ndc1 is not solely a component of NPCs but also part of spindle pole bodies (SPBs), the NE-embedded microtubule-organizing centers (Araki et al., 2006; Chial *et al*., 1998; Winey et al., 1993). Ndc1 is involved in insertion of both complexes into the intact NE during interphase (Chial *et al*., 1998; Jaspersen and Ghosh, 2012; West et al., 1998; Winey *et al*., 1993). In yeast cells, Ndc1 is essential. Depletion causes cell cycle arrest, mislocalization of soluble nucleoporins to cytosolic foci and also affects the morphology of the NPC in the NE (Madrid *et al*., 2006; Onischenko *et al*., 2009; Winey *et al*., 1993).

Ndc1 interacts with the non-essential linker nucleoporins Nup53 and Nup59 thereby bridging between the core of the NPC and the NE, conserved in metazoan (Eisenhardt et al., 2014; Mansfeld *et al*., 2006; Onischenko *et al*., 2009). Both Nup53 and Nup59 additionally contain C-terminal amphipathic helices thereby exhibiting membrane binding and deformation activity (Eisenhardt *et al*., 2014; Marelli et al., 2001; Patel and Rexach, 2008; Vollmer et al., 2012). Furthermore, curvature-sensing proteins are recruited to the NE e.g. via ArfGAP1 lipid packing sensor (ALPS) motifs (Drin et al., 2007; Vanni et al., 2013), thought to stabilize the thermodynamically unfavoured curved membrane state. Nup120 and Nup133, conserved members of the Y-shaped Nup84 complex (Kelley et al., 2015; Lutzmann et al., 2002; Stuwe et al., 2015), contain ALPS motifs in exposed unstructured loops of their β-propeller domains capable of binding and stabilizing positive membrane curvature via membrane insertion (Doucet et al., 2015; Doucet et al., 2010; Drin and Antonny, 2010; Drin *et al*., 2007; Kim et al., 2014; Nordeen et al., 2020). Deletion of *NUP120* or *NUP133* causes NPC clustering in the NE underlying the proposed function of ALPS motives in NPC biogenesis (Li et al., 1995; Pemberton et al., 1995; Siniossoglou et al., 1996). In addition to the ALPS motif-mediated targeting to the NE, vertebrate Nup107-Nup160 complex can directly bind the membrane nucleoporin POM121 via the N-terminal β-propeller region of Nup160 (Nup120 in lower eukaryotes), (Mitchell et al., 2010). Furthermore, the Nup107-Nup160 complex can be recruited to membranes via interaction with the nuclear basket protein Nup153 which itself can bind the INM via an amphipathic helix (Vollmer et al., 2015).

In this study we show that in *Chaetomium thermophilum* both Nup120 and Nup133 can be recruited to membranes independently of their ALPS domains via direct protein-protein interactions with Ndc1. In addition, we focus on a so far uncharacterized C-terminal region of the essential yeast Ndc1 protein which is dispensable for viability and therefore not further being suggested to having any crucial functions (Lau *et al*., 2004). We show that this region contains a previously unknown conserved amphipathic motif and elucidate its interrelationship within the nucleoporin network at the NE in the context of interphase NPC biogenesis in *Saccharomyces cerevisiae*.

## MATERIALS AND METHODS

### Media, yeast strains, plasmids and genetic methods

Media preparation and microbiological techniques were performed according standard protocols (Ausubel et al., 1992; Guthrie and Fink, 1991). *Saccharomyces cerevisiae* strains used in this study were listed in Table S1. Genetic manipulations were carried out using a homologous recombination-based approach and verified by colony PCR (Gueldener et al., 2002; Janke et al., 2004). Cloning and plasmid propagation were performed according to standard procedures using *E. coli* DH5α. Site-directed mutagenesis was performed using a PCR-based method (Pfirrmann et al., 2013). Plasmids used in this study were listed in Table S2. Yeast and *E. coli* expression plasmids for proteins from *Chaetomium thermophilum* were generated using gene amplification from a cDNA library (Amlacher et al., 2011). Yeast expression plasmids based on YEplac112 additionally possess a *LEU2* marker under control of the truncated *leu2d* promoter increasing plasmid copy number if corresponding cells grow on media lacking leucine (Erhart and Hollenberg, 1983). Proteins expressed from the YEplac112-based plasmids are N-terminally tagged with two IgG binding units of Protein A from *Staphylococcus aureus* (Rigaut et al., 1999) followed by a TEV protease recognition sequence (ProtA-TEV). All plasmids generated were verified by sequencing.

### Protein expression and purification

Full length soluble nucleoporins from *Chaetomium thermophilum* carrying an N-terminal ProtA-TEV tag were expressed in *S. cerevisiae* under control of the *GAL1-10* promoter and affinity-purified using IgG beads. Briefly, cells were grown overnight in raffinose medium (SRC-Leu) to an OD_600_ of 2 prior to dilution with galactose medium (2 x YPG). After 6 hours of protein expression, cells were harvested, resuspended in lysis buffer (20 mM HEPES pH 7.6, 150 mM NaCl, 50 mM KAc, 2 mM Mg(Ac)_2_, 5 % (w/v) glycerol, 0.1 % (w/v) NP-40 and protease inhibitor mix (Sigma-Aldrich)) and lysed mechanically using glass beads. After centrifugation at 39,000 x g at 4 °C for 20 min the supernatant was incubated with 0.5 ml of pre-equilibrated IgG-Sepharose slurry (GE Healthcare) at 4 °C for 1 h. After three washing steps bound proteins were subjected to TEV protease cleavage for 1 h at 16 °C (TEV protease expressed and purified in house). Eluted proteins were collected and further used for in *vitro* binding assays. Cells expressing the membrane nucleoporins *Ct*Ndc1 and *Ct*Pom152 were expressed as described earlier but resuspended in a differently composed lysis buffer (20 mM Tris pH 7.6, 150 mM NaCl, 50 mM KAc, 2 mM Mg(Ac)_2_, 5 % (w/v) glycerol, 2 % (w/v) Triton X-100, 2 mM DTT and protease inhibitor mix) prior to immobilization on IgG beads for subsequent *in vitro* binding assays.

*Ct*Ndc1, *Ct*Ndc1 truncations and SCL1/BC08 (Lorenz et al., 2015) for reconstitution in GUVs were expressed in *E. coli* BL21 codon plus (DE3) cells, (EMD Millipore) from a modified pET28a plasmid containing an N-terminal MISTIC (membrane-integrating sequence for translation of integral membrane protein constructs) fragment (13 kDa) from *Bacillus subtilis* allowing high yield expression of membrane proteins in *E. coli* (Roosild et al., 2005) followed by a thrombin protease cleavage site. Both proteins were additionally expressed as C-terminal 6 x His fusion proteins enabling Ni-based affinity purification as described elsewhere (Alves et al., 2017). Briefly, purification was done in buffer containing 1 % (w/v) cetyltrimethylammonium bromide (CTAB) on Ni-NTA magnetic beads (EMD) followed by dialysis against PBS containing 1 % (w/v) CTAB and 1 mM EDTA. MISTIC was finally cleaved off using thrombin protease (GE Healthcare).

Truncated versions of *Ct*Nup120 and *Ct*Nup133 used for *in vitro* binding assays were expressed and affinity-purified from *E. coli* BL21 codon plus (DE3) cells. Briefly, corresponding proteins were expressed from modified pPROEX-1 (Invitrogen)-based plasmids as N-terminal 6 x His-TEV fusion proteins. Cells were grown to an OD_600_ of 0.5 prior to IPTG-mediated induction (1 mM) of expression at 30 °C for 3 h. Cells were harvested and resuspended in lysis buffer (20 mM HEPES pH 7.5, 200 mM NaCl, 1 mM DTT and protease inhibitor mix). Lysis was performed using a high-pressure cavitation homogenizer (Microfluidizer 110L, Microfluidics) followed by centrifugation at 39,000 x g at 4 °C for 20 min. Precleared lysates were incubated with Ni-NTA beads (Macherey-Nagel) for 1 h at 4 °C. After extensive washing in buffer containing 10 mM imidazole pH 8.0 bound proteins were eluted in buffer containing 200 mM imidazole.

### Fluorescence microscopy

Fluorescence microscopy was done with cells expressing GFP-tagged *Sc*Nup49 serving as a marker for NPC integrity in the NE. Chromatin was stained using Hoechst 33258 dye. Briefly, cells were grown overnight in selective media (plus 2 x adenine) to mid-exponential growth phase. 1 ml of the cell suspension was harvested and resuspended in 800 µl buffered glucose (100 mM HEPES pH 7.7, 2 % (w/v) glucose). The chromatin stain Hoechst 33258 (Sigma-Aldrich) was added (10 µg/ml) and the cells afterwards incubated in the dark for 3 min. Cells were fixed with formaldehyde (3 % (w/v)) for 5 min, washed two times and resuspended in 25 µl buffered glucose. Fluorescence microscopy was performed using a microscope (Imager Z1; Carl Zeiss) with a 100 x, NA 1.4 Plan-Apochromat oil immersion objective lens (Carl Zeiss). Pictures were acquired with a CCD camera (AxioCamM Rm; Carl Zeiss) and the AxioVision SE64 Rel. 4.9.1 imaging software (Carl Zeiss).

### Transmission electron microscopy

Yeast cells were grown overnight in liquid media to mid-exponential growth phase and resuspended in fixation buffer (0.2 M PIPES pH 6.8, 0.2 M sorbitol, 2 mM MgCl_2_, 2 mM CaCl_2_, 2 % (w/v) formaldehyde and 2 % (w/v) glutaraldehyde) for 10 at RT and 50 min at 4 °C. After washing in 0,1 M phosphate-citrate buffer, pH 5.8 the cells were resuspended in the same buffer containing 0.25 mg/ml Zymolyase 20T (Seikagaku Corp., Tokyo) and incubated for 2 h at 30 °C. The cells were washed three times in 0.1 M sodium acetate buffer pH 6.1 and post-fixed with 2 % (w/v) osmium tetroxide/1.5 % (w/v) K_4_[Fe(CN)_6_] in 0.1 M sodium phosphate buffer pH 7.4. After contrasting en bloc with 0.5 % (w/v) uranyl acetate overnight and dehydrating using a graded dilution series of ethanol the cells were embedded into glycid ether 100-based resin. Ultrathin sections were prepared with a Reichert ultracut S ultramicrotome (Leica). Thin sections were collected on formvar-coated grids and contrasted with both uranyl acetate and lead citrate. Specimens were visualized using an electron microscope (EM 10 CR; Carl Zeiss, Oberkochen) equipped with a 1K CCD camera (Tröndle Restlichtverstärkersysteme, Moorenweis) at an acceleration voltage of 60 KV. Imaging was done by the software ImageSP (SYSPROG).

### Lipid analysis

Yeast cells were grown overnight in corresponding media, harvested in mid-exponential growth phase and resuspended in 50 mM HEPES pH 7.4 containing 0.8 mg/ml Zymolyase 100T (Carl Roth). Cells were mechanically lysed using glass beads. Lipidomics analyses were performed by a shotgun approach as recently described (Papagiannidis et al., 2021). Lipid extracts were prepared in presence of internal lipid standards via acidic Bligh-Dyer lipid extraction (Bligh and Dyer, 1959). The used master mix of lipid internal standards contained 50 pmol d_7_-PC mix (15:0/18:1-d_7_, Avanti Polar Lipids), 25 pmol PI (17:0/20:4, Avanti Polar Lipids), 50 pmol PE and 10 pmol PS (14:1/14:1, 20:1/20:1, 22:1/22:1, semi-synthesized as described in (Ozbalci et al., 2013)), 15 pmol PA (PA 17:0/20:4, Avanti Polar Lipids), 10 pmol PG (14:1/14:1, 20:1/20:1, 22:1/22:1), semi-synthesized as described in (Ozbalci *et al*., 2013)), 40 pmol DAG (17:0/17:0, Larodan), 40 pmol TAG (D_7_-TAG-Mix, LM-6000/D5-TAG 17:0,17:1,17:1, Avanti Polar Lipids), 10 pmol t-Cer (18:0, Avanti Polar Lipids) and 50 pmol ergosteryl ester (15:0 and 19:0). The lipids in the organic phase were dried under a gentle stream of nitrogen at 37 °C. Lipids were resuspended in 10 mM methanolic ammonium acetate and transferred to 96-well plates (Eppendorf Twintec 96). Mass spectrometry was performed on a Sciex QTRAP 6500+ mass spectrometer, equipped with chip-based (HD-D ESI Chip; Advion Biosciences) nano-electrospray infusion, and ionization (TriVersa NanoMate; Advion Biosciences) as described in (Ozbalci *et al*., 2013). The following precursor ion (PREC) or neutral loss (NL) scanning modes were used: +PREC184 (PC), +PREC282 (t-Cer), +NL141 (PE), +NL185 (PS), +NL277 (PI), +NL189 (PG), +NL115 (PA), +NL77 (ergosterol), +PREC379 (ergosteryl ester). Mass spectrometric analysis of ergosterol was performed applying a one-step chemical derivatization to ergosterol acetate in the presence of 300 pmol (first and second replicate) or 100 pmol (third and fourth replicate) of the internal standard (22E)-Stigmasta-5,7,22-trien-3-beta-ol (Sigma-Aldrich, R202967) using 100 μl acetic anhydride/chloroform (1:12 v/v) overnight under argon atmosphere according to (Ejsing et al., 2009). Data evaluation was done using LipidView (Sciex) and ShinyLipids (in house developed software).

### Reconstitution of proteins into GUVs

Prior to reconstitution of the detergent-solubilized membrane proteins *Ct*Ndc1 and SCL1/BC08 in GUVs the purified proteins were labeled using a succinimidyl ester of Alexa Fluor 488 dissolved in 200 mM NaHCO_3_ pH 8.4 containing 1 % (w/v) CTAB. Detergent removal and subsequent formation of proteoliposomes were performed via gel filtration after incubation of the detergent-solubilized and labeled proteins with a lipid mixture mimicking the composition of the NE (Eisenhardt *et al*., 2014; Lorenz *et al*., 2015). Briefly, the protein-lipid mixture was loaded on a Sephadex G50 fine filled Econo chromatography column equilibrated in sucrose buffer (10 mM HEPES pH 7.5, 250 mM sucrose, 50 mM KCl and 2.5 mM MgCl_2_), pelleted and resuspended in 20 mM HEPES pH 7.4, 100 mM KCl and 1 mM DTT. GUVs containing either *Ct*Ndc1 or SCL1/BC08 were reconstituted via electroformation after vacuum drying onto 5 x 5 mm platinum gauzes as described elsewhere (Lorenz *et al*., 2015).

Successful reconstitution of *Ct*Ndc1 and SCL1/BC08 into GUVs was monitored by light emission between 500-545 nm after argon laser-mediated excitation at 488 nm using a LSM 710 microscope (Carl Zeiss) with a 40 x, NA 1.2 C-Apochromat water immersion objective lens. Recruitment of Alexa546-labeled nucleoporins to either *Ct*Ndc1-GUVs or SCL1/BC08-GUVs was monitored by collection of emitted light between 570-625 nm after excitation at 546 nm.

### Liposome flotation assays

Liposome generation and flotation was performed as described in (Vollmer *et al*., 2015). In short, *E. coli* polar lipids (Avanti Polar Lipids) dissolved in chloroform and supplemented with 0.2 mol % octadecyl rhodamin B chloride (R18, Thermo Fisher) were vacuum dried on a rotary evaporator, dissolved as liposomes in PBS by freeze/thawing cycles and extruded by passages through Nuclepore track-etched membranes (Whatman) with defined pore sizes using an Avanti Mini-Extruder to generate small unilamellar liposomes of defined sizes. For liposome flotations proteins purified from either pGEX-4T3 or pET28a-based plasmids (6 μM) were mixed 1:1 with liposomes (5 mg/ml) and floated for 2h at 55,000 rpm in a TLS-55 rotor (Beckman) at 25 °C through a sucrose gradient. Binding efficiency was determined by Western Blot analysis using a ImageQuant LAS-4000 system (Fuji) and the AIDA software, comparing band intensities of start materials with floated liposome fractions.

### Miscellaneous

SDS-PAGE and Western Blot analysis were performed as described elsewhere (Laemmli, 1970; Towbin et al., 1979). PageRuler^TM^ Unstained (Thermo Scientific), PageRuler^TM^ Prestained Protein Ladder (Thermo Scientific) and peqGOLD Protein-Marker I Unstained (VWR) were used as protein markers. Immunodetection of ProteinA (ProtA)-tagged proteins was performed using Peroxidase Anti-Peroxidase Soluble Complex antibody (Sigma-Aldrich) in a 1:3,000 dilution. Brilliant Blue G-Colloidal Concentrate Electrophoresis Reagent (Sigma-Aldrich) served as Coomassie stain.

## RESULTS

### Nup120 and Nup133 can interact with Ndc1 *in vitro* and are recruited to Ndc1 containing GUVs

The Nup84 complex (Nup107-Nup160 complex in vertebrates) forms a scaffold coating and presumably stabilizing the high curvature of the nuclear pore membrane (see Introduction). To find out whether integral nuclear pore membrane proteins play a role in the recruitment and anchoring of the Nup84 complex to the NPC in fungi, we performed *in vitro* binding studies using nucleoporins from the thermophilic fungus *Chaetomium thermophilum*. For this purpose, we expressed *Ct*Ndc1 and *Ct*Pom152 in yeast, affinity-purified both membrane proteins in detergent and tested for interaction with members of the *Ct*Nup84 complex. This analysis revealed a previously unknown direct interaction of either *Ct*Nup120 or *Ct*Nup133 with the integral membrane nucleoporin *Ct*Ndc1, whereas *Ct*Nup84 as another component of the Nup84 subcomplex (Thierbach et al., 2013) did not bind to *Ct*Ndc1 (Fig. 1A). In a negative control, *Ct*Pom152 did not show such an interaction with *Ct*Nup120 or *Ct*Nup133 (Fig. 1A).

**Figure 1.**
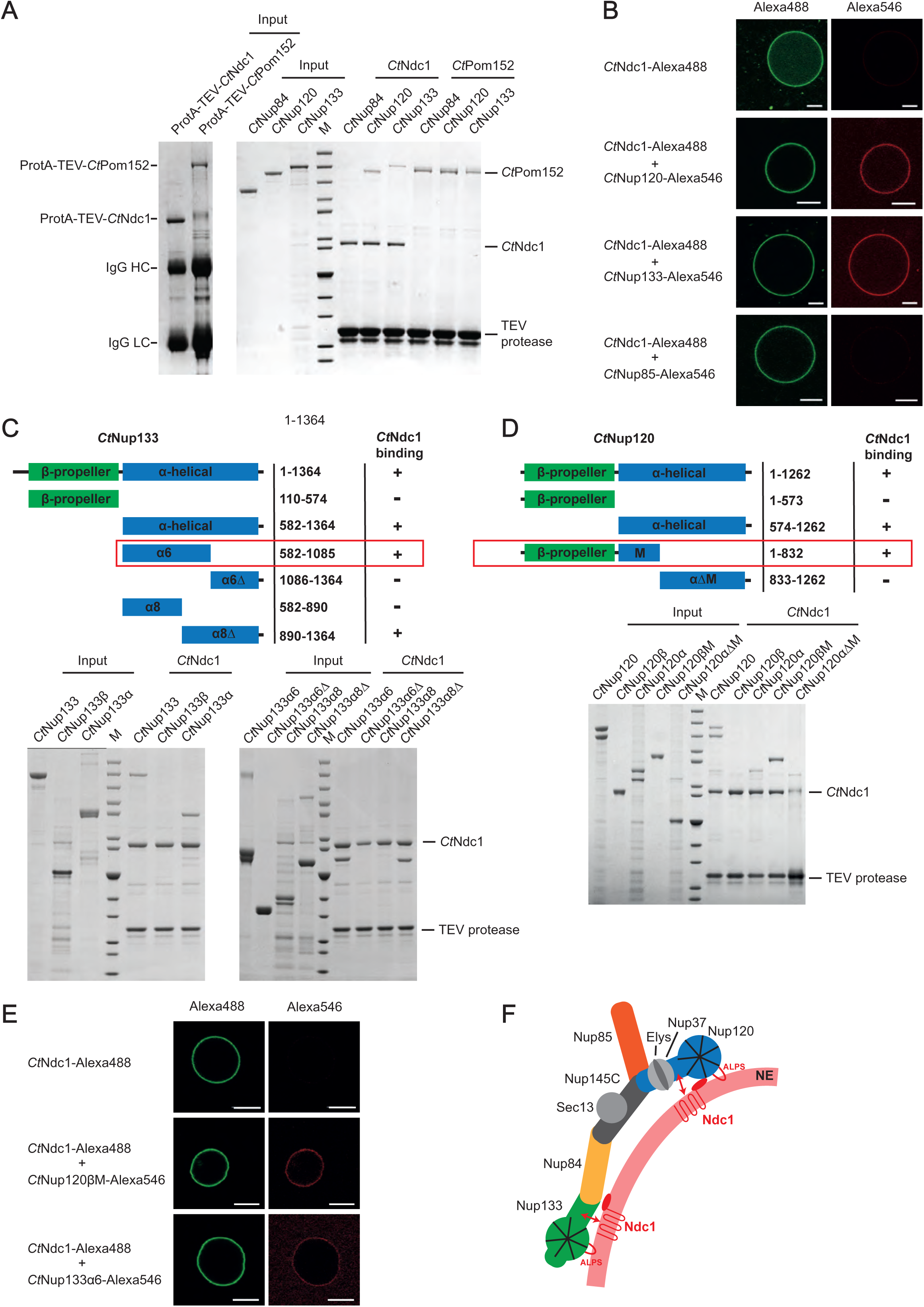
*Chaetomium* Nup120 and Nup133 interact with Ndc1 and are recruited to Ndc1-containing GUVs. (A) *In vitro* binding of nucleoporins of the *Ct*Nup84 complex to *Ct*Ndc1 and *Ct*Pom152. ProteinA (ProtA)-tagged *Ct*Ndc1 and *Ct*Pom152 were expressed in *Saccharomyces cerevisiae* and immobilized on IgG beads. The beads were incubated with affinity-purified nucleoporins of the *Ct*Nup84 complex. Proteins bound to IgG were eluted with TEV protease cleaving downstream the ProtA affinity tag of *Ct*Ndc1 and *Ct*Pom152. Input and elution fractions were analysed via SDS-PAGE and Coomassie staining. (B) Alexa Fluor 488-labeled *Ct*Ndc1 was reconstituted into giant unilamellar vesicles (GUVs). Ndc1 GUVs were monitored by collection of emitted light between 500-545 nm after argon laser-mediated excitation at 488 nm (left panels) whereas recruitment of Alexa Fluor 546-labeled *Ct*Nup120, *Ct*Nup133 or *Ct*Nup85 (red) to *Ct*Ndc1-GUVs (green) was analysed by confocal microscopy. Scale bars, 10 µm. (C) *In vitro* binding of truncated versions of *Ct*Nup133 and (D) *Ct*Nup120 purified in *E. coli* BL21 (DE3) to ProtA-tagged *Ct*Ndc1 immobilized on IgG beads as described earlier. (E) Recruitment of either Alexa546-labeled *Ct*Nup120βM or *Ct*Nup133α6 (red) to *Ct*Ndc1-GUVs (green) was analysed as above (middle panels). Scale bars, 10 µm. (F) Model of recruitment and stabilization of the *Ct*Nup84 complex and *Ct*Ndc1 repectively, to sites of NPC assembly in the NE. The *Ct*Nup84 complex binds directly to the NE via exposed ALPS domains in the beta propeller domains of both *Ct*Nup120 and *Ct*Nup133 and also interacts via alpha-helical domains of *Ct*Nup120 and *Ct*Nup133 with the membrane nucleoporin *Ct*Ndc1.

To reveal this Ndc1-Nup84 complex interaction not only in a micellar detergent system, but in a real membrane environment, we reconstituted purified *Ct*Ndc1 (Fig. S1A) into giant unilamellar vesicles (GUVs). This physiological-like reconstitution assay showed *Ct*Nup120 and *Ct*Nup133 (Fig. S1B) recruitment to *ct*Ndc1 embedded in GUV membranes, whereas *Ct*Nup85 was still not bound (Fig. 1B). Moreover, *Ct*Nup120 and *Ct*Nup133 did not associate with GUVs containing the unrelated inner nuclear membrane protein BC08/SCL1 (Fig. 1C). Although possessing ALPS domains, membrane association of both *Ct*Nup120 and *Ct*Nup133 in absence of Ndc1 was probably hindered by the low curvature of the GUV membranes. This is similar to the vertebrate Y-complex, which does not bind to GUVs (Vollmer *et al*., 2015). Mapping of the protein-protein interaction sites revealed that sequences in the C-terminal α-helical domains of *Ct*Nup120 and *Ct*Nup133 mediated *Ct*Ndc1 binding, whereas the β-propeller domains did not bind to *Ct*Ndc1 (Fig. 1C, D). Further fine-mapping indicated that the truncated α-helical domain of *Ct*Nup133 (*Ct*Nup133α6) was still able to bind to *Ct*Ndc1 (Fig. 1C). Since the truncated α-helical domain of *Ct*Nup120 (*Ct*Nup120 (574-832)) was insoluble, we succeeded expressing it together when still attached to the β-propeller domain (*Ct*Nup120βM), which finally revealed strong binding of *Ct*Nup120βM to *Ct*Ndc1 (Fig. 1D). Accordingly, purified *Ct*Nup133α6 and *Ct*Nup120βM (Fig. S1D) were efficiently recruited to *Ct*Ndc1-GUVs (Fig. 1E), but not to SCL1/BC08 containing GUVs (Fig. S1E). To find out which parts of *Ct*Ndc1 bind to *Ct*Nup120 and *Ct*Nup133, we separated *Ct*Ndc1 N-terminal transmembrane (Fig. S1F (I)) and C-terminal soluble domains (Fig. S1F (II)). This revealed that only the *Ct*Ndc1 N-terminal transmembrane part was sufficient to recruit *Ct*Nup120 or *Ct*Nup133 to the GUV membrane (Fig. S1F (I)).

Together, these data suggest that the *Ct*Nup84 complex, besides its ALPS-mediated interaction with the NE (see Introduction), has a second way to attach to the nuclear pore membrane, namely via direct interaction of the Ndc1 transmembrane domain with either *Ct*Nup120 or *Ct*Nup133. In this way, the integral part of the pore membrane protein Ndc1 could function in local Nup84 complex attachment at the fused and curved double nuclear membrane, which is the site where NPCs are inserted (Fig. 1F).

### Overexpression of Ndc1 induces ER membrane expansions with pore-like holes

In the course of *GAL*-promoter induced ProtA-*Ct*Ndc1 expression in yeast, we noticed the formation of extensive extranuclear membranes, tubules and cisternae, which were not observed in the control strain expressing only ProtA (Fig. 2A-B). These abnormal membranous structures may be derived from the rough ER, which is continuous with the outer nuclear membrane and cortical ER. Similar membrane proliferations were observed earlier upon overexpression of GFP-*Hs*NDC1 or GFP-*Hs*POM121 in human HeLa cells (Volkova et al., 2011). Serial ultrathin sections of fixed yeast cells overexpressing ProtA-*Ct*Ndc1 revealed that these unusual extranuclear membrane proliferations exhibited pore-like structures with diameters similar to the diameter of NPCs within the nuclear membrane (Fig. 2C). Overexpression of Ndc1 from *Saccharomyces cerevisiae* (ProtA-*Sc*Ndc1) induced similar membrane morphologies (Fig. 2D). However, overexpression of the ER membrane protein 3-hydroxy-3-methylglutaryl coenzyme A reductase (ProtA-*Sc*Hmg1), which is known to induce formation of karmellae, i.e. membrane stacks associated with the outer nuclear membrane (Wright et al., 1988), clearly lacked these pore-like structures (Fig. 2E).

**Figure 2.**
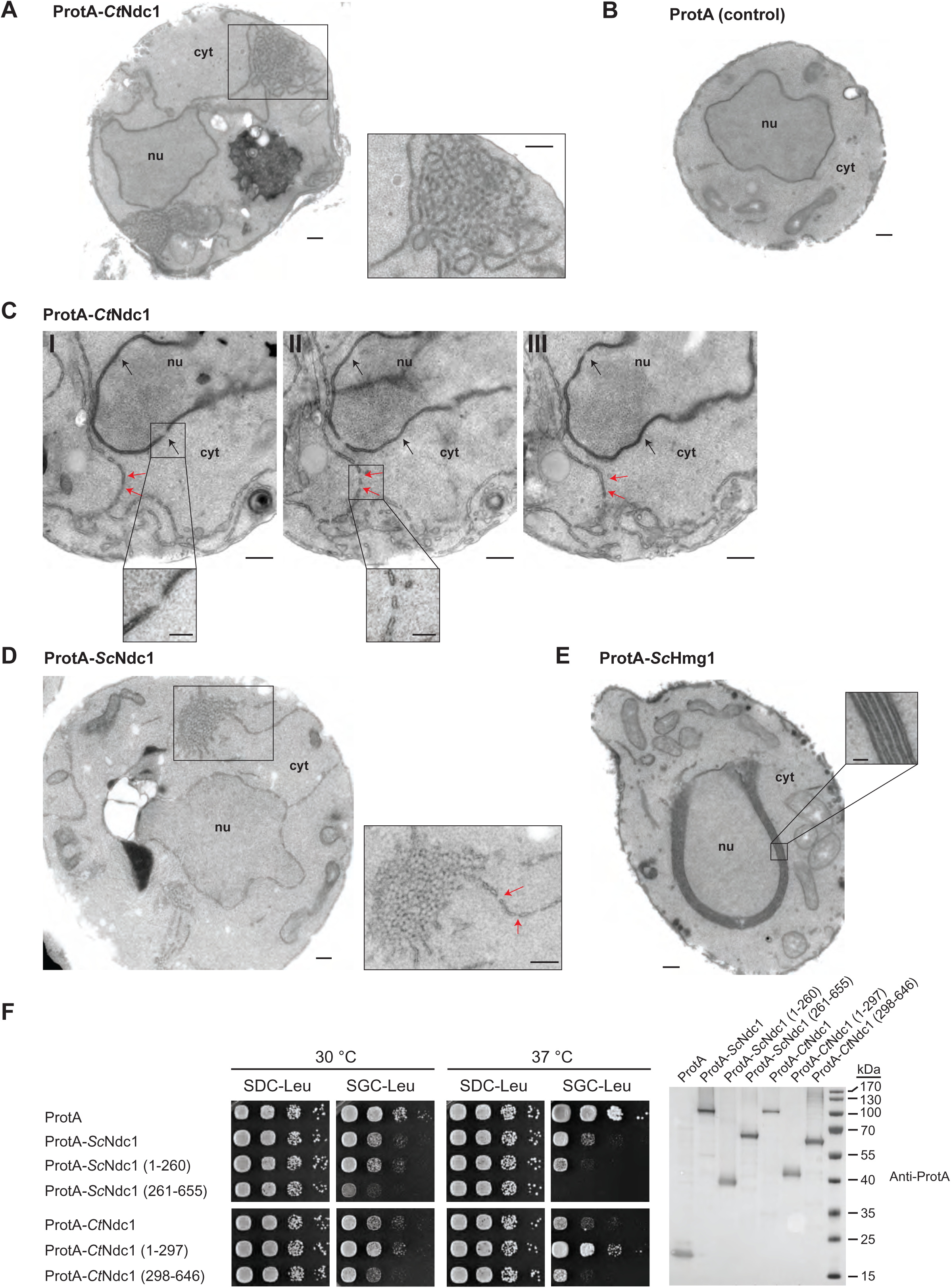
Overexpression of Ndc1 in budding yeast induces ER membrane expansion with pore-like structures. (A, B) Transmission electron microscopy of a wild-type yeast strain transformed with ProtA-*Ct*Ndc1 (A) or as control ProtA tag alone (B) stained with osmium tetroxide plus potassium ferrocyanide. Magnifications (right panel show the extranuclear membrane assemblies). Bars, 250 nm. (C) Serial sections (approximate thickness of 80 nm) of a cell overexpressing ProtA-*Ct*Ndc1. Black arrows mark nuclear pore complexes visible in the NE. Red arrows mark local ring-like openings in the extranuclear membranes. Bars, 250 nm. Magnifications show a nuclear pore in the NE and membrane openings observed upon overexpression of ProtA-*Ct*Ndc1 (lower panels; bars, 100 nm). (D) Transmission electron microscopy of a yeast cell overexpressing ProtA-*Sc*Ndc1. Cytoplasmic membrane assemblies with local membrane discontinuities, marked with arrows, are magnified; bars, 250 nm). (E) Transmission electron microscopy of a yeast cell overexpressing ProtA-tagged 3-hydroxy-3-methylglutaryl-coenzymeA reductase 1 (ProtA-*Sc*Hmg1). Karmellae observed upon ProtA-*Sc*Hmg1 expression were magnified (bar, 250 nm and 50 nm for the magnified panel). (F) Growth tests with yeast cells overexpressing ProtA-tagged full length Ndc1 and Ndc1 truncations from both *C. thermophilum* and *S. cerevisiae*, ProtA-*Sc*Hmg1 or ProtA alone using *LEU2* as selection marker. 10-fold serial dilutions of corresponding overnight cultures were spotted onto either glucose (expression suppressing) or galactose (expression inducing) containing plates lacking leucine (SDC-Leu or SGC-Leu) and incubated at 30 °C or 37 °C for up to three days. For expression control whole cell lysates were prepared from overnight cultures and analysed by Western blotting.

Consistent with the observed dramatic membrane proliferation phenotype upon overexpression of either *Ct*Ndc1 or *Sc*Ndc1 in yeast, these cells exhibited significant growth defects both at 30 °C and 37 °C when compared to wild-type yeast cells expressing ProtA alone (Fig. 2F). To find out, which of the Ndc1 domains is responsible for the observed toxic effects, we overexpressed either the C-terminal (soluble cytoplasmic/nucleoplasmic) or N-terminal (trans-membrane) part of Ndc1 from either *S. cerevisiae* or *C. thermophilum* in yeast. In both cases, the dominant growth defect could be predominantly attributed to the corresponding Ndc1 C-domains, and the growth inhibition was even more severe than observed for the overexpression of the full-length Ndc1 proteins (Fig. 2F, Fig. S2C). Overexpression of ProtA-*Sc*Hmg1 was also highly toxic to yeast cells (Fig. S2C). Altogether, these findings showed that Ndc1 overexpression induced a strong membrane proliferation phenotype, yielding extranuclear membranes showing pore-like perforations with a size similar to the NPC diameter.

### The cytoplasmic/nucleoplasmic C-terminal domain of Ndc1 contains a conserved amphipathic motif

To identify the toxicity-inducing motif in the Ndc1 C-domain, we generated a serious of Ndc1 deletion constructs. In the past, several *Sc*Ndc1 truncations have been already constructed, e.g. by removing a large part from the non-essential C-terminus (Fig. S2A, *Sc*Ndc1 Δ368-466), but also other parts upstream and downstream of residues 368-466, which were suggested to be crucial for *Sc*Ndc1 function and localization (e.g. for interaction with the SPB components *Sc*Nbp1 and *Sc*Mps3 (Chen et al., 2014; Lau *et al*., 2004). Prompted by the observation that overexpression of Ndc1 can induce membrane proliferation, we searched for a possible amphipathic motif in the *Sc*Ndc1 C-terminal domain by HELIQUEST (Gautier et al., 2008), which indeed detected such a sequence between residues 448-465 (Fig. 3A). The primary structure of this predicted amphipathic sequence shows a low conservation, even among closely related *Saccharomycetales*, but its amphipathic character is strongly conserved (Fig. 3A). We cloned this predicted amphipathic helix (AH) from yeast Ndc1 and inserted it between N-terminal ProtA and C-terminal eGFP. After expression and affinity-purification of this ProtA-AH_Ndc1_-eGFP construct, we tested whether it can bind to an artificial membrane employing a liposome flotation assay (Fig. 3B). This revealed that ProtA-AH_Ndc1_-eGFP can efficiently bind to liposomes, with a preference for smaller and highly curved liposomes, which is typical for the well-characterized ALPS (ArfGAP1 lipid packing sensor) motifs (Drin *et al*., 2007; Vanni *et al*., 2013). When disturbing the hydrophobic face of the putative amphipathic helix by changing Ndc1 L461>D, this mutation strongly reduced liposome binding (Fig. 3B). To analyse whether the Ndc1-AH alone is capable of inducing membrane abnormalities upon overexpression in yeast, we analysed the intracellular location of ProtA-AH_Ndc1_-eGFP constructs by fluorescence microscopy in yeast. This revealed cytoplasmic foci and patches, often seen close to either the plasma membrane (likely to be cortical ER) or nuclear membrane, whereas in the case of the mutant Ndc1-AH only a continuous cytoplasmic staining was seen (Fig. 3C). Similar distribution patterns were also observed for cells expressing the Ndc1-AH when part of the entire *Sc*Ndc1 C-terminus (Fig. S3A, eGFP-*Sc*Ndc1 (261-655) and eGFP-*Sc*Ndc1 L461D (261-655). As anticipated, the C-terminal domains of the employed higher eukaryotic Ndc1 homologs from *Xenopus laevis* or mouse were also capable of binding to liposomes, whereas the purified C-terminal part of *Xl*Nup98 serving as negative control was not recruited (Fig. S3B).

**Figure 3.**
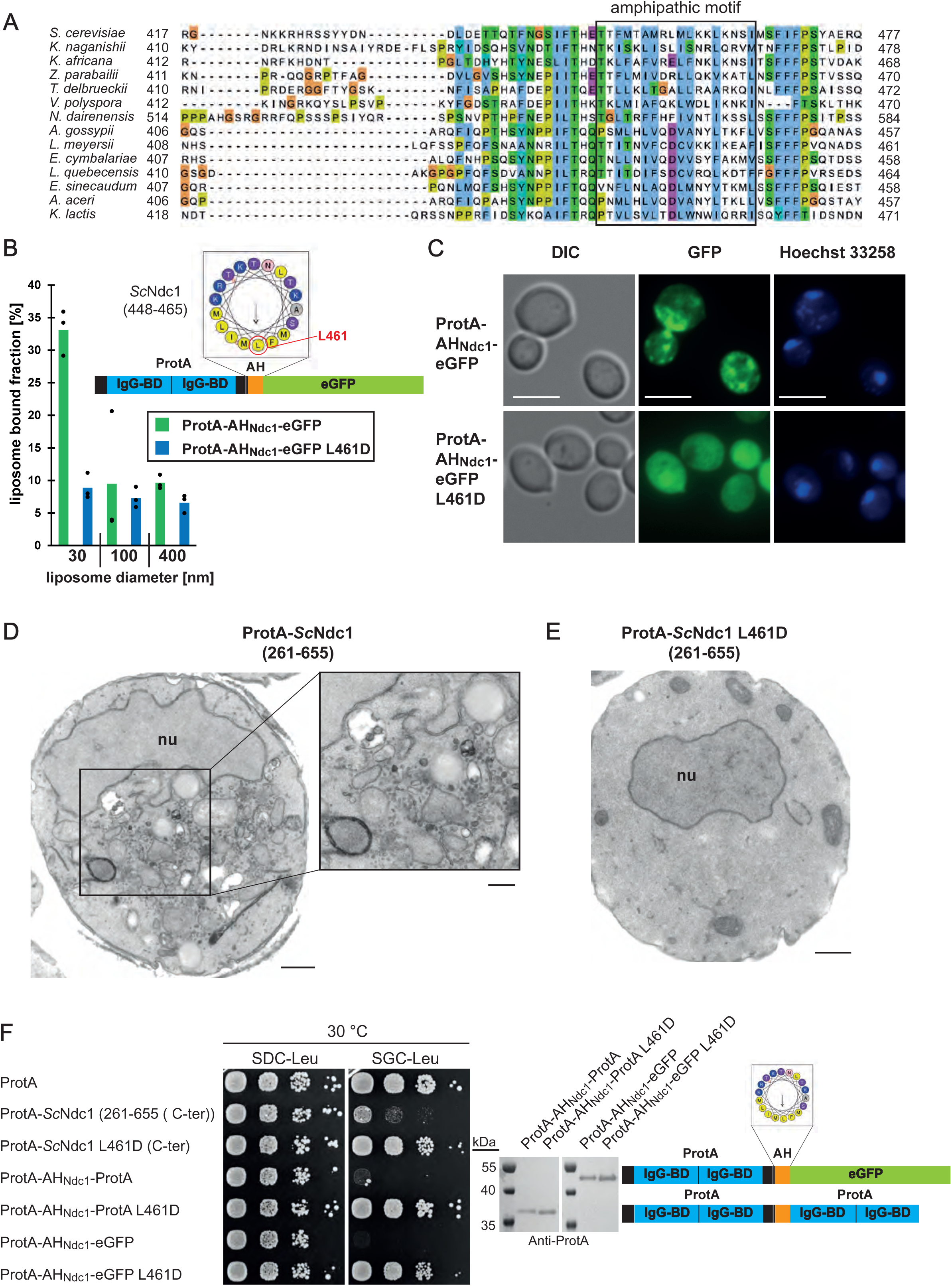
The C-terminus of Ndc1 contains a conserved amphipathic sequence motif. (A) Multiple sequence alignment of a C-terminal region of Ndc1 homologues from different fungal species. The alignment was done using the ClustalW (Larkin et al., 2007) and displayed in Jalview (Waterhouse et al., 2009). The putative amphipathic regions in all sequences were detected using HELIQUEST (Gautier *et al*., 2008). (B) *In vitro* liposome binding assay with the isolated amphipathic helix (ProtA-AH_Ndc1_-eGFP) and the corresponding L461D mutation using 30 nm, 100 nm and 400 nm liposomes. Columns represent the mean of three independent experiments, individual data points are indicated. (C) Fluorescence microscopy of cells overexpressing the isolated amphipathic helix (ProtA-AH_Ndc1_-eGFP) and the corresponding L461D mutation. Chromatin is stained with Hoechst 33258. Bars, 5 µm. (D) Electron microscopy of a yeast cell expressing the C-terminal part of *Sc*Ndc1 (ProtA-*Sc*Ndc1 (261-655)); bar, 500 nm. The cytoplasm is additionally magnified (right panel; bar, 250 nm). (E) Electron microscopic visualization of yeast cell expressing ProtA-*Sc*Ndc1 (261-655) carrying the point mutation L461D. Bar, 500 nm. (F) Growth and expression were analysed as in Fig. 2F using yeast cells overexpressing the artificial model constructs ProtA-AH_Ndc1_-ProtA and ProtA-AH_Ndc1_-eGFP. The domain structures of both artificial constructs were schematically illustrated.

To reveal the specific membrane alterations induced by the toxic Ndc1 amphipathic motif in more detail, we performed transmission electron microscopy of cells overexpressing ProtA-*Sc*Ndc1 (261-655). Surprisingly, many cells accumulated extra cytoplasmic membrane structures, in particular clusters of small vesicles (Fig. 3D). Similar results were observed in yeast overexpressing the ALPS-domain-containing α-synuclein, which induced curved membrane structures including a large number of small vesicles (Outeiro and Lindquist, 2003; Pranke et al., 2011). Cells expressing the Ndc1 C-terminus carrying the mutated AH at L461>D (ProtA-*Sc*Ndc1 L461D (261-655)), however, exhibited a normal membrane organization (Fig. 3E) like cells expressing ProtA alone (Fig. 2B).

To prove that the Ndc1-AH is responsible for the toxicity initially observed when overexpressing the complete C-terminus of Ndc1 in yeast cells (Fig. 2F), we generated minimal constructs, ProtA-AH_Ndc1_-eGFP and ProtA-AH_Ndc1_-ProtA, either intact or carrying the L461D mutation, and induced them *in vivo* via the strong *GAL1-10* promoter, which yielded a similar growth pattern like observed for the longer constructs (Fig. 3F). Furthermore, we generated chimeric *Sc*Ndc1 constructs composed of ProtA-*Sc*Ndc1 (261-655), but carrying instead of its own amphipathic helix (residues 448-465) the amphipathic motifs from other Ndc1 orthologs including human, or the well-characterized amphipathic helices of *Sc*Nup53 and *Sc*Nup59. Except for the AH from *Sc*Nup59, all the other AHs induced a toxic phenotype in yeast (Fig. S3C). Replacing the Ndc1-AH with an 18 amino acid long stretch of alternating alanine-serine residues, called (AS)_9_, or deleting the complete AH (ΔAH) rescued the growth defect (Fig. S3C). Moreover, only L461D or L461E point mutations, destroying the hydrophobic character of the Ndc1-AH, but not a conserved L461I mutation, could inactivate the toxicity upon overproduction (Fig. S3D).

The growth defects observed upon overexpression of either ProtA-*Sc*Ndc1 (448-655) or solely by ProtA-AH_Ndc1_-ProtA were also accompanied by altered cellular lipid profiles (Fig. S4A). In particular, we observed in the case of the intact Ndc1-AH overproduction a massive reduction of the cellular phospholipids phosphatidylethanolamine (PE) and phosphatidylserine (PS) whereas the amount of storage lipids increased in comparison to control cells. The Ndc1 L461D mutation, however, reversed this altered lipid pattern to normal levels profiles as observed in the control strain expressing only ProtA (Fig. S4A). The cellular total lipid content was further increased upon overexpression of the Ndc1-AH-containing constructs, which was also noticed for ProtA-*Sc*Hmg1, which might be due to the extensive membrane proliferation and/or lipid storage in these strains (Fig. S4B-C).

Since also overexpression of ProtA-*Sc*Nup53 or ProtA-*Sc*Nup59 was toxic to cells, which, like in the case of the Ndc1-AH, was caused by the corresponding C-terminal amphipathic motifs (Fig. S5A), we wondered why ProtA-*Sc*Nup53 overproduction triggered formation of inner nuclear membrane abnormalities (Fig. S5B; see also (Marelli *et al*., 2001)), whereas Ndc1-AH caused cytoplasmic membrane abnormalities (see Fig. 3D). We suspected that *Sc*Nup53 may exert its toxic effect in the nucleus due to NLS-mediated nuclear import (Lusk et al., 2002). Hence, we fused the *Sc*Ndc1 C-domain to the NLS of SV40 large T antigen (Kalderon et al., 1984). As anticipated, this construct, which was still toxic for the cells, induced extensive intranuclear membrane proliferations (Fig. S5D-E).

Taken together, a transferable and conserved amphipathic motif in the C-terminus of *Sc*Ndc1, similar to related motifs in functionally interacting nucleoporins such as *Sc*Nup53, exerts a dominant-negative effect on intracellular membrane organization, which may have its primary cause in affecting membrane curvatures (see Discussion).

### Ndc1’s essential function can be suppressed by deleting the Nup53 amphipathic helix

Prompted by the finding that Ndc1 and Nup53/Nup59 amphipathic motifs may the nuclear membrane curvature at the NPC insertion sites in a coordinated fashion, we wondered whether this is also manifested in genetic relationship, for instance in form of a synthetic lethal interaction. However, we discovered the opposite, namely that the lethal phenotype of a complete *NDC1* gene disruption was rescued by further deleting the non-essential *NUP53* gene, which quite unexpectedly allowed the *ndc1*Δ *nup53*Δ double disruption strain to regain growth (Fig. 4A). In contrast, the chromosomal *nup59*Δ deletion did not rescue the *ndc1*Δ cells (Fig. 4A), whereas a *ndc1*Δ *nup53*Δ *nup59*Δ triple knock-out strain was still viable (Fig. 4A). Notably, also the a *pom34*Δ deletion could restore growth of the lethal *ndc1*Δ strain (Fig. 4A). Thus, the absence of the non-essential integral pore membrane protein Pom34 or lack of the linker Nup Nup53, which via its C-terminal amphipathic helix interacts with the nuclear membrane (Eisenhardt *et al*., 2014; Marelli et al., 1998; Patel and Rexach, 2008; Vollmer *et al*., 2012), rescues the lethal phenotype of the *ndc1*Δ null yeast strain.

**Figure 4:**
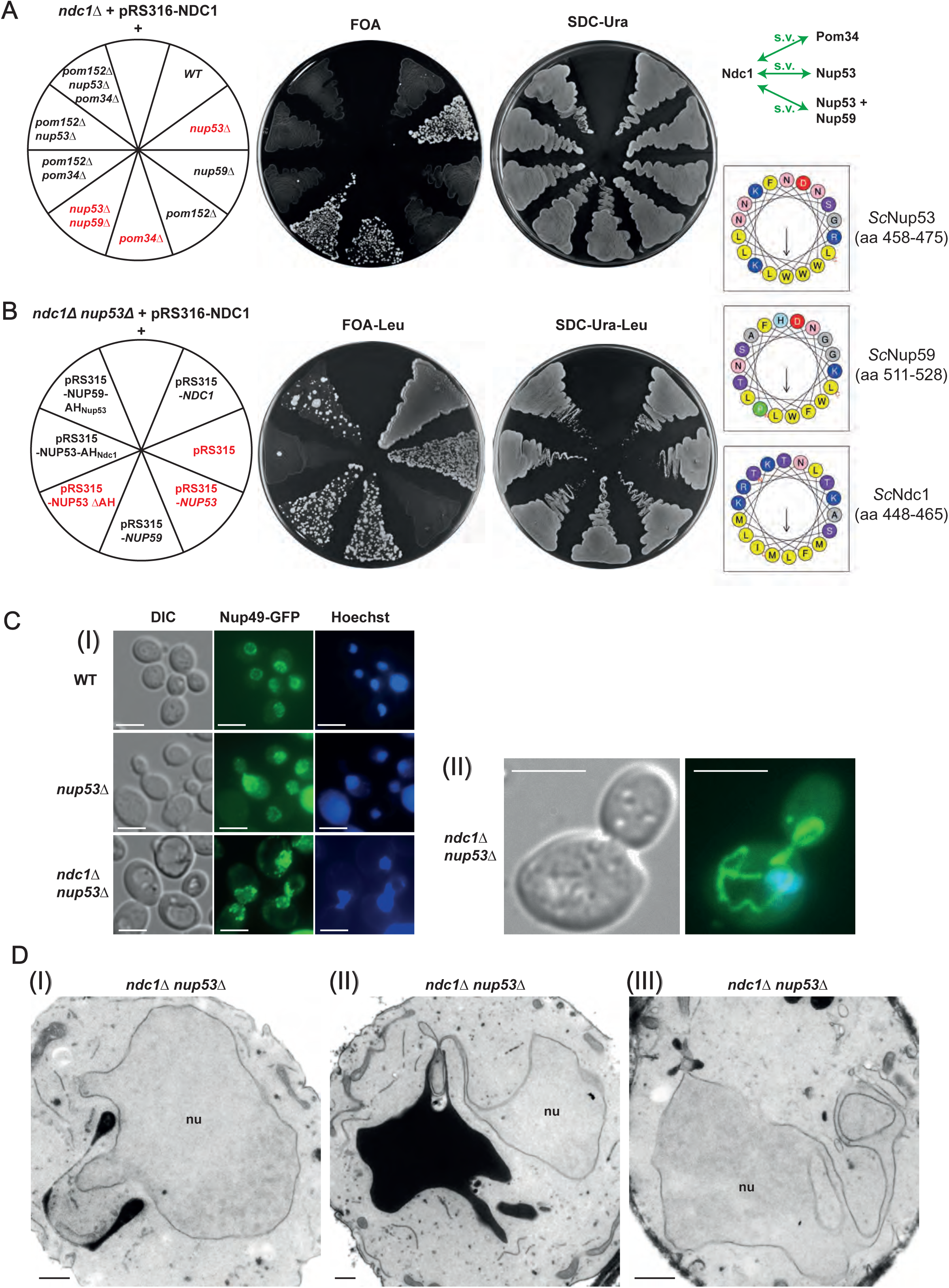
*Sc*Ndc1 genetically interacts with the amphipathic helix of *Sc*Nup53. (A) Nup53 and Pom34 interact genetically with Ndc1. The used yeast strains were all endogenously deleted in *NDC1* and expressed *Sc*Ndc1 from a centromeric *URA3* marker-containing plasmid under control of its native promoter (pRS316-*ScNDC1*). Loss of pRS316-*ScNDC1* was forced by spreading out on plates containing 5-fluoroorotic acid (FOA). Growth on SDC-Ura plates served as control. Green arrows in the illustration mark the observed positive genetic interactions resulting in synthetic viability (s.v.) of corresponding double/triple deletion strains. (B) Ndc1 shuffle strains deleted in *NUP53* and expressing different proteins, as indicated, from pRS315, a centromeric *LEU2* marker-containing plasmid, were spread on FOA-Leu plates forcing loss of pRS316-*ScNDC1.* Growth on SDC-Ura-Leu plates served as control selecting for presence of both plasmids. The amphipathic motifs present in the expressed proteins were illustrated as helical wheel projections. (C) (I) Fluorescence microscopy of yeast strains, as indicated, all expressing the eGFP-tagged nucleoporin *Sc*Nup49 as NPC marker. Hoechst 33258 served as chromatin stain. (II) Yeast cells deleted in both *NDC1* and *NUP53* were additionally magnified visualizing nuclear membrane protrusions into the cytoplasm. Bars, 5 µm. (D) Electron microscopic visualization of yeast cells (I-III) deleted in both *NDC1* and *NUP53*. Bars, 500 nm.

To find out whether the *nup53*Δ dependent suppression of *ndc1*Δ is dependent on the Nup53 C-terminal amphipathic helix (AH), we combined *nup53* ΔAH and *ndc1*Δ alleles, which however still allowed the cells to grow (Fig. 4B). In contrast, replacing Nup53’s C-terminal amphipathic helix by the AH_Ndc1_ abolished the suppression phenotype and *ndc1*Δ cells were no longer viable (Fig. 4B, Nup53-AH_Ndc1_). On the other hand, a Nup59-AH_Nup53_ construct enabled the *ndc1*Δ suppression phenotype (Fig. 4B) indicating that Nup59 performs (an) unique role(s) at the NE independent of its C-terminal AH.

To further characterize this unusual *ndc1*Δ *nup53*Δ suppressor strain, we analysed the subcellular distribution of the nucleoporin marker Nup49-eGFP. Whereas wild-type and *nup53*Δ cells exhibited predominantly round-shaped nuclei, the Nup49-eGFP staining observed in *ndc1*Δ *nup53*Δ cells revealed nuclei with irregular shapes, often with an isthmus-like restriction or with extensive nuclear membrane protrusions, which reached deeply into the cytoplasm and were devoid of chromatin (Fig. 4C I-II). Electron microscopic analysis of *ndc1*Δ *nup53*Δ cells revealed these NE abnormalities and extensions in greater detail (Fig. 4D I-III), which were also observed in *ndc1*Δ *nup53*Δ *nup59*Δ cells (Fig. S6B I) or *ndc1*Δ *pom34*Δ cells (Fig. S6B II). Interestingly, highly related NE abnormalities were reported in earlier studies, in one case upon deletion of *NEM1* or *SPO7*, which encode a phosphatase complex controlling phospholipid biosynthesis and nuclear membrane growth (Siniossoglou et al., 1998), and in the other case upon overexpression of the NE-associated Esc1, which is involved in nuclear basket organization (Hattier et al., 2007; Lewis et al., 2007).

Consistent with a disturbed nuclear membrane biogenesis, the deletion strains *ndc1*Δ *nup53*Δ, *ndc1*Δ *nup53*Δ *nup59*Δ and *ndc1*Δ *pom34*Δ were found to be sensitive to the membrane-fluidizing drug benzyl alcohol (BA) (Fig. S6A). In addition, the microtubule-depolymerizing drug benomyl diminished cell growth of corresponding strains, which may be also a sign that SPB biogenesis is impaired in the absence of Ndc1 (Fig. S6A). Finally, overexpression of Faa3, a long chain fatty acyl-CoA synthetase, rescued the growth defect of *ndc1*Δ *nup53*Δ cells upon exposure to BA (Fig. S6C). A similar effect of Faa3-induced recovery was observed for the BA-sensitive *mps2*Δ *pom152*Δ strain, which also showed abnormalities in NE morphology (Friederichs et al., 2011).

Thus, the essential Ndc1 becomes dispensable for cell growth in yeast if the amphipathic helix of Nup53 is deleted, but such cells exhibit an abnormal NE morphology, which may be attributed to an altered lipid composition with a nuclear membrane biogenesis (see Discussion).

### The amphipathic helix of Ndc1 becomes crucial in a *nup59*Δ strain

Next, we studied the function of the Ndc1-AH within the nucleoporin network, which is part of the NPC inner ring scaffold (Fig. 5A). Since both Nup53 and Nup59 via their C-terminal AH motifs insert into the nuclear pore membrane and both interact with Ndc1, an overlapping function of these highly homologous linker Nups has been suggested (Marelli *et al*., 2001; Onischenko *et al*., 2009; Patel and Rexach, 2008; Vollmer *et al*., 2012). On the other hand, Nup53 and Nup59 also fulfil unique roles, which among other things is indicated by the observation that the *nup59*Δ, but not *nup53*Δ deletion strain, is lethal when combined with *pom34*Δ or *pom152*Δ knock-out alleles (Marelli *et al*., 1998; Miao et al., 2006; Onischenko *et al*., 2009).

**Figure 5.**
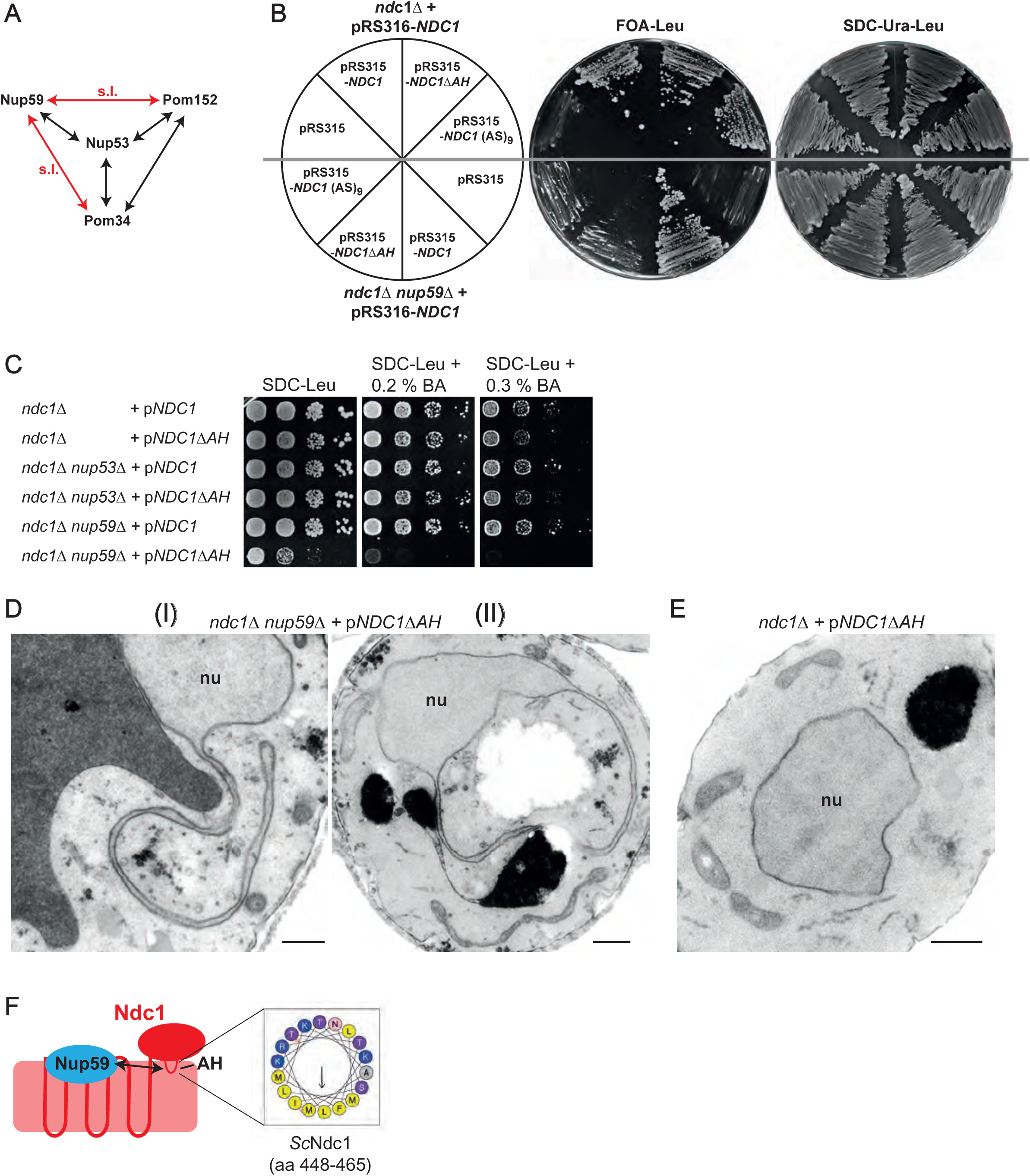
The Ndc1-AH becomes crucial if Nup59 is lacking. (A) Illustration of known negative genetic interactions within the interaction network of nucleoporins of the inner ring and the linker nucleoporins Nup53 and Nup59. Black arrows mark negative genetic interactions upon deletion of corresponding genes. Red arrows mark negative genetic interactions resulting in synthetic lethality (s.l.) of corresponding double deletion strains. (B) Cells from *Sc*Ndc1 shuffle strains deleted in *NUP59* and additionally expressing *Sc*Ndc1 variants encoded by *LEU2* marker-containing pRS315 plasmids, as indicated, were spread onto FOA-Leu plates forcing plasmid loss of pRS316-*ScNDC1*. Growth on SDC-Ura-Leu served as control. (C) Growth of *ndc1*Δ, ndc1Δ *nup*53Δ and *ndc1*Δ *nup59*Δ cells expressing wild-type *Sc*Ndc1 or *Sc*Ndc1ΔAH from pRS315-based plasmids was monitored on leucine-lacking plates containing different amounts of the membrane-fluidizing drug benzyl alcohol (BA). (D (I-II)) and (E) Electron microscopic visualization of either (D (I-II)) *ndc1*Δ *nup59*Δ or (E) *ndc1*Δ cells expressing *Sc*Ndc1ΔAH from a pRS315-based plasmid. Bars, 500 nm. (F) Graphical illustration of the observed interaction between the amphipathic motif of *Sc*Ndc1 and *Sc*Nup59.

Based on this genetic data, we expressed Ndc1ΔAH as well as a related construct, Ndc1 (AS)_9_, carrying instead of the AH a tandem alanine-serine linker sequence, in the *nup59*Δ *ndc1*Δ double disruption strain (Fig. 5B). Strikingly, corresponding cells displayed a slow-growth phenotype compared to *NUP59* cells expressing Ndc1 lacking the amphipathic motif (Fig. 5B).

To further investigate these opposite effects exerted by Ndc1-AH in the single *ndc1*Δ versus *ndc1*Δ *nup59*Δ double disruption strains, we incubated cells with the membrane-fluidizing drug benzyl alcohol (BA) during a dot spot growth analysis (Fig. 5C). This revealed that the already strong growth inhibition of the *nup59*Δ *ndc1*Δ cells expressing Ndc1ΔAH was converted into a complete growth inhibition by addition of 0.2 % BA (Fig. 5C). However, ndc*1*Δ or *ndc1*Δ *nup53*Δ cells complemented by Ndc1ΔAH did not show such a BA-induced growth inhibition (Fig. 5C).

To detect changes in the intracellular membrane organization of cells expressing Ndc1ΔAH but lacking Nup59, we performed electron microscopy (Fig. 5D). This revealed long nuclear membrane protrusions reaching deeply into the cytoplasm, similar to what has been already observed in the case of *ndc1*Δ *nup53*Δ (Fig. 4D, II), *ndc1*Δ *nup53*Δ *nup59*Δ (Fig. S6B, I) or *ndc1*Δ *pom34*Δ cells (Fig. S6B, II). In contrast, these membrane abnormalities were not observed in *ndc1*Δ cells complemented by Ndc1ΔAH (Fig. 5E).

Together, this data show that the amphipathic motifs in Ndc1 and Nup59 exhibit a specific genetic interaction (Fig. 5F) by which this linker Nup can perform a unique role at the NE not related to the Ndc1-AH function.

## DISCUSSION

In this study, we describe new functional motifs of the nuclear membrane protein Ndc1, which enlighten its complex physical and genetic interactions within the NE-associated nucleoporin network. The N-terminal part of Ndc1 mediates direct contact to the Y-shaped Nup84 complex, whereas a short amphipathic helix that functionally interacts with related motifs in the inner pore ring linker Nups, Nup53 and Nup59, directly binds the pore membrane.

Conserved ALPS domains present in the β-propeller domains of both Nup120/Nup160 and Nup133 (Drin *et al*., 2007; Kim *et al*., 2014; Kosinski et al., 2016; von Appen et al., 2015) presumably contribute to the targeting the Nup84 complex to the curved nuclear pore membrane. Using *in vitro* binding assays, we found that a direct interaction between the C-terminal α-helical domains of both *Ct*Nup120 and *Ct*Nup133 and the N-terminus of Ndc1, pointing to an additional mechanism of how the Nup84 complex is recruited to the pore membrane. Interestingly, in *C. elegans* embryos Nup160 as a Y-complex component is partially mislocalized in Ndc1-depleted cells (Mauro et al., 2022). It is thus conceivable that the Y-complex in addition to its ALPS motifs requires direct interactions to transmembrane nucleoporins as Ndc1 for NPC localization. Notably, in vertebrates the Y-complex can bind the integral pore membrane protein POM121 (Mitchell *et al*., 2010). Here, it remains open whether vertebrate NDC1 could in addition to POM121 serve as recruitment point for the Y-complex.

Membrane interaction motifs have been identified in several nucleoporins with crucial roles in NPC assembly and function (Hamed and Antonin, 2021). Here, we identify a membrane binding motif in a transmembrane nucleoporin. This motif is localized within the C-terminal half of Ndc1 and preferentially binds to highly curved membranes, and hence behaves like a classical ALPS (Vanni et al., 2013), despite possessing also charged lysine and arginine residues in its polar face (Fig. 3B). This ALPS motif might be involved in targeting Ndc1 to the curved pore membrane and/or contribute to the curved shape of the pore membrane. In vertebrates, the N-terminus of Ndc1 is essential but not sufficient for NPC assembly (Eisenhardt et al., 2014). This hinted towards an essential function of the C-terminus of Ndc1 during NPC biogenesis, probably involving the membrane binding role described here. In the case of yeast Ndc1, which has a dual role in NPC and SPB insertion, it is assumed that it acts as a mobile and dynamic membrane protein, which can be recruited to assembly sites of both NPCs and SPBs.

Surprisingly, the essential yeast Ndc1 is dispensable for cell growth if Nup53 is absent (Fig. 4A), and this suppression phenotype depends on the C-terminal amphipathic helix of Nup53 (Fig. 4B). In *in vitro* NPC reconstitution assays the N-terminal transmembrane regions of vertebrates Ndc1 are only required if Nup53 has a functional C-terminal amphipathic helix (Eisenhardt *et al*., 2014) pointing to a conserved function of the Nup53-Ndc1 interaction. Ndc1 is not essential in *Aspergillus nidulans*, but this fungus also lacks obvious Nup53 orthologs (Liu et al., 2009; Osmani et al., 2006). The phylum *Amoebozoa* lacks both Ndc1 and Nup53 homologs or possesses Nup53 homologs with only poor sequence homology at the C-terminus (Neumann *et al*., 2010). This and our results are in line with the idea that the N-terminus of Ndc1 controls or modulates the membrane bending activity of Nup53 (Eisenhardt *et al*., 2014).

The observed dramatic changes in the NE morphology observed for *NDC1*-deleted cells that on top lack Nup53, Pom34 or both, Nup53 and Nup59 may be due to alterations in NE membrane homeostasis tightly linked to NPC biogenesis. The Apq12-Brr6-Brl1 complex, which is conserved in organisms undergoing closed mitosis (Tamm et al., 2011), transiently interacts with early NPC intermediates (Lone et al., 2015; Zhang et al., 2021; Zhang et al., 2018). The mechanistic role of this complex is not clear but a direct or indirect role in promoting phosphatidic acid accumulation at the curved membranes was suggested (Zhang *et al*., 2021). This, in turn might stabilize or even induce membrane curvature via the conical shaped phosphatidic acid (Zhukovsky et al., 2019). As Ndc1 interacts with Brl1 (Zhang *et al*., 2018), it is conceivable that absence of Ndc1 the Apq12-Brr6-Brl1 complex is not properly recruited to NPC assembly sites.

The three viable *NDC1*-deletion strains (*ndc1*Δ *nup53*Δ, *ndc1*Δ *nup53*Δ *nup59*Δ and *ndc1*Δ *pom34*Δ) are sensitive towards the membrane-fluidizing drug benzyl alcohol (Fig. S6A) suggesting defects in NE lipid homeostasis. Overexpression of long chain fatty acid synthetase Faa3 rescued the BA sensitivity of *ndc1*Δ *nup53*Δ cells, similar as observed for the *pom152*Δ *mps2*Δ strain (Friederichs *et al*., 2011). Presumably, increasing *de novo* lipid synthesis of long chain fatty acid-containing membrane lipids stabilizes the curvature of the pore membrane and by this counteract the increased membrane fluidity of cells caused by disturbances of the lipid homeostasis system.

In contrast to *NUP53*, deletion of *NUP59* did not suppress the lethal *ndc1*Δ disruption. We initially speculated that the differences in the amino acid composition of their C-terminal amphipathic helices are responsible for this discrepancy. However, expressing the chimeric protein Nup59-AH_Nup53_ consisting of Nup59 and the amphipathic motif of Nup53 still allowed suppressive growth of *ndc1*Δ cells. Thus, both orthologs possess distinct functions, which is consistent with previous results (Marelli *et al*., 1998; Miao *et al*., 2006; Onischenko *et al*., 2009). This is further supported by our genetic analyses of the amphipathic helix in the C-terminus of Ndc1 and Nup59 (Fig. 5B). Cells deleted in *NUP59* expressing Ndc1ΔAH display a severe growth defect and sensitivity towards BA. Nup59 is involved in proper localization of Ndc1 to the NE overlapping in its function with the Pom152-Pom34 subcomplex (Onischenko *et al*., 2009). Presumably, the Ndc1-AH is involved in correct positioning of Ndc1 at the NPCs for subsequent integration into the nucleoporin network. Ndc1ΔAH might not be targeted properly to the NE if Nup59 is absent and therefore corresponding cells behave similar to those lacking Ndc1.

In conclusion, our work expands the repertoire of the NE-interacting motifs by a previously unknown amphipathic motif in the C-terminus of Ndc1. It reveals that membrane interaction motifs in different nucleoporins, here Ndc1 and Nup53, influence each other. Whether this also includes the ALPS of the Y-complex or of other Nups is an interesting avenue for future research. Finally, our data indicate a so far unrecognized targeting mechanism of the Nup84 complex to the NE by direct interactions with Ndc1, possibly enhancing the efficiency of the ALPS domain-mediated targeting to the curved NE. Future studies on this question may help to clarify whether this interaction is conserved in vertebrates, where POM121 does bind to the Y-complex.

## SUPPLEMENTARY DATA

**Supplemental Figure 1.**
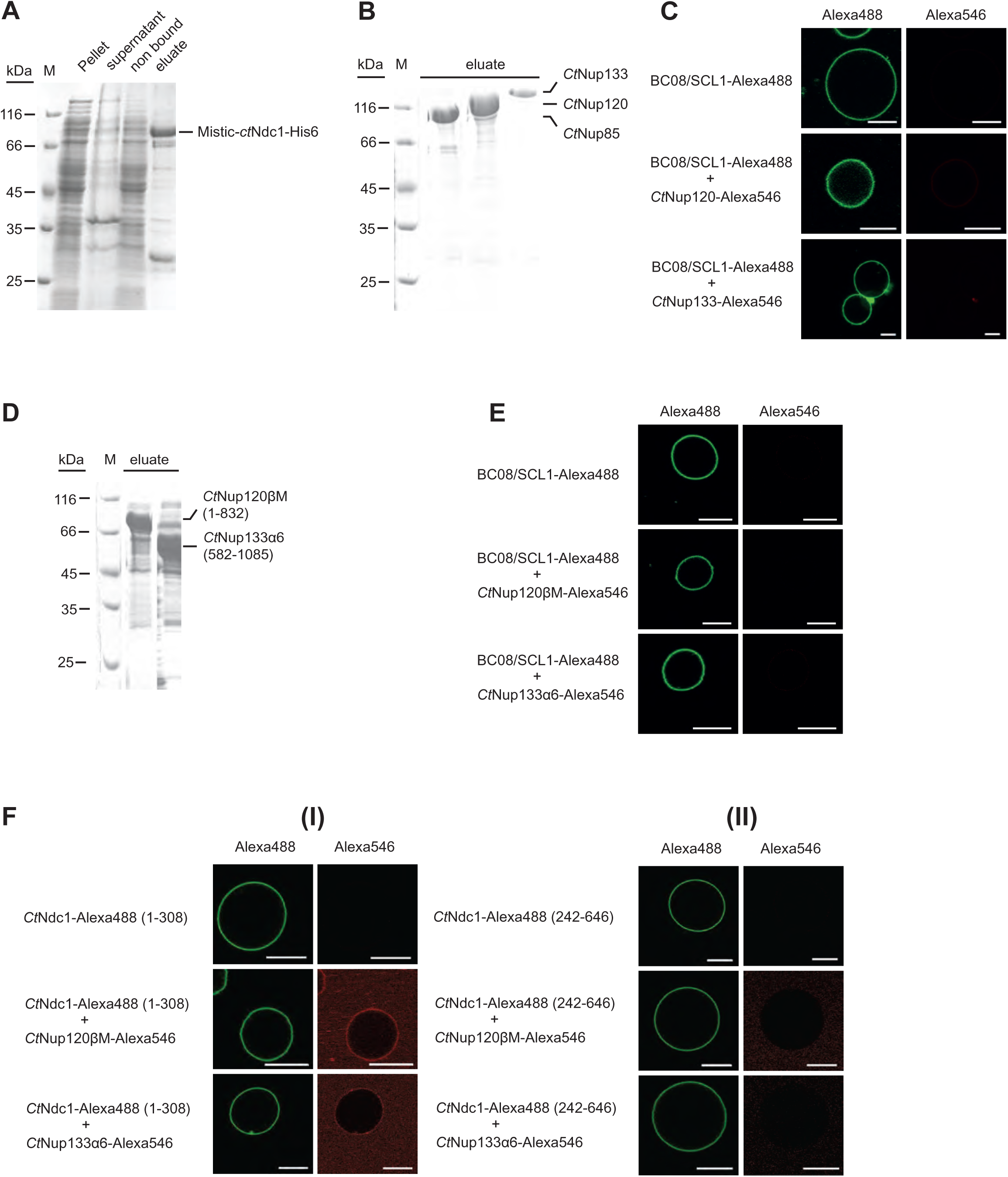
(A) *Ct*Ndc1 used for reconstitution in GUVs was Ni-NTA-purified from *E. coli* BL21 (DE3). *Ct*Ndc1 was additionally N-terminally fused to MISTIC, a *B. subtilis* protein allowing high-level expression. Elution was done by imidazole treatment. Prior to GUV reconstitution, the MISTIC tag was cleaved off using thrombin protease. The different fractions obtained during purification were analysed via SDS-PAGE and Coomassie staining. (B) ProteinA-tagged *Ct*Nup85, *Ct*Nup120 and *Ct*Nup133 were expressed in *Saccharomyces cerevisiae* under control of the *GAL1-10* promoter and purified using IgG beads. Purified proteins were cleaved off the beads using TEV protease. Elution fractions were subjected to SDS-PAGE and Coomassie staining. (C) SCL1/BC08 was expressed and purified in *E. coli* BL21 (DE3), as described for *Ct*Ndc1, Alexa488-labeled and reconstituted into GUVs. SCL1/BC08-GUVs were incubated with either Alexa546-labeled Nup120 or Nup133. Binding was monitored as described earlier. Bars, 10 µm. (D) Expression and purification of *Ct*Nup120βM consisting of an α-helical extension of the β-propeller domain of *Ct*Nup120 and *Ct*Nup133α6, an alpha-helical stretch in the C-terminal region of *Ct*Nup133. Elution fractions were subjected to SDS PAGE and Coomassie staining. (E) SCL1/BC08-GUVs were incubated with either Alexa546-labeled Nup120βM or Nup133α6. Binding was monitored as described earlier. Bars, 10 µm. (F) (I) N-(1-308) or (II) C-terminal (242-646) fragments of *Ct*Ndc1, expressed, purified a labeled as above, were reconstituted into GUVs and incubated with Alexa546-labeled Nup120βM or Nup133α6. The *Ct*Ndc1 (242-646) fragment additionally harbored the sixth transmembrane helix of *Ct*Ndc1 to ensure proper reconstitution into GUV membranes.

**Supplemental Figure 2.**
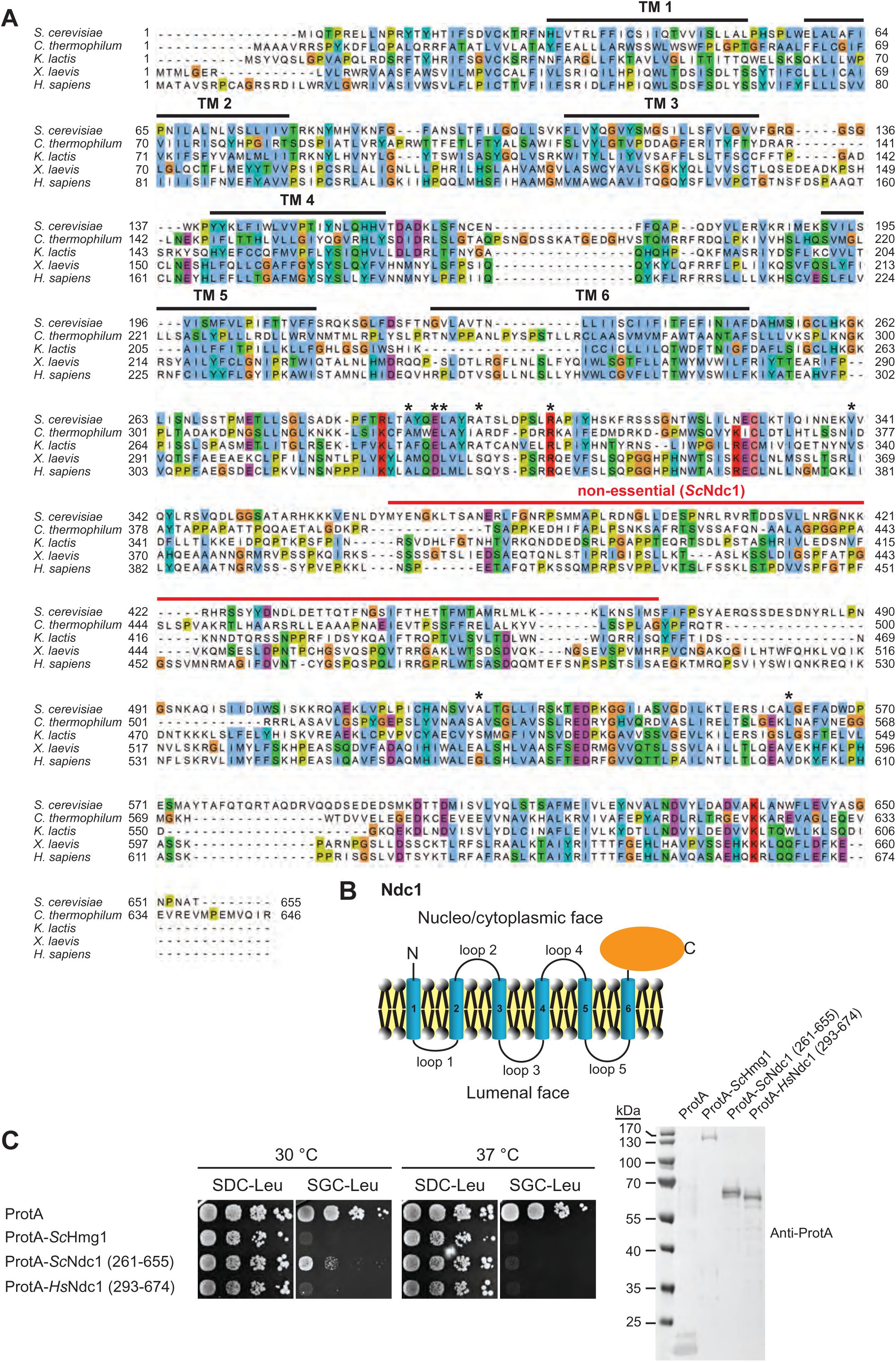
(A) Multiple sequence alignment of Ndc1 homologs from *S. cerevisiae* (CAA86624.1), *C. thermophilum* (AEL00695.1), *K. lactis* (QEU62154.1), *X. laevis* (ABA39292.1) and *H. sapiens (*AAZ73087.1) using the ClustalW (Larkin *et al*., 2007) and displayed in Jalview (Waterhouse *et al*., 2009). The transmembrane domains (TM) were predicted using TMHMM (Krogh et al., 2001) and are marked with bold black lines. The red line marks the region in *Sc*Ndc1 (368-466) discovered as not being essential for *Sc*Ndc1 function (Lau *et al*., 2004). Asterisks mark amino acid positions in the C-terminus of *Sc*Ndc1 involved in interactions with the SPB components *Sc*Mps3 and *Sc*Nbp1 respectively (Chen *et al*., 2014). (B) Topology of Ndc1. Ndc1 consists of six predicted transmembrane domains connected via short loops. Both the N-terminal end and the C-terminal part of Ndc1 are exposed to the cytoplasm and nucleoplasm (Stavru *et al*., 2006). (C) Growth and expression were analysed as described earlier using yeast cells expressing both ProtA-tagged Ndc1 homologs from *S. cerevisiae and H. sapiens* and ProtA-tagged *Sc*Hmg1 under control of the *GAL1-10* promoter from *LEU2* marker-containing plasmids.

**Supplemental Figure 3.**
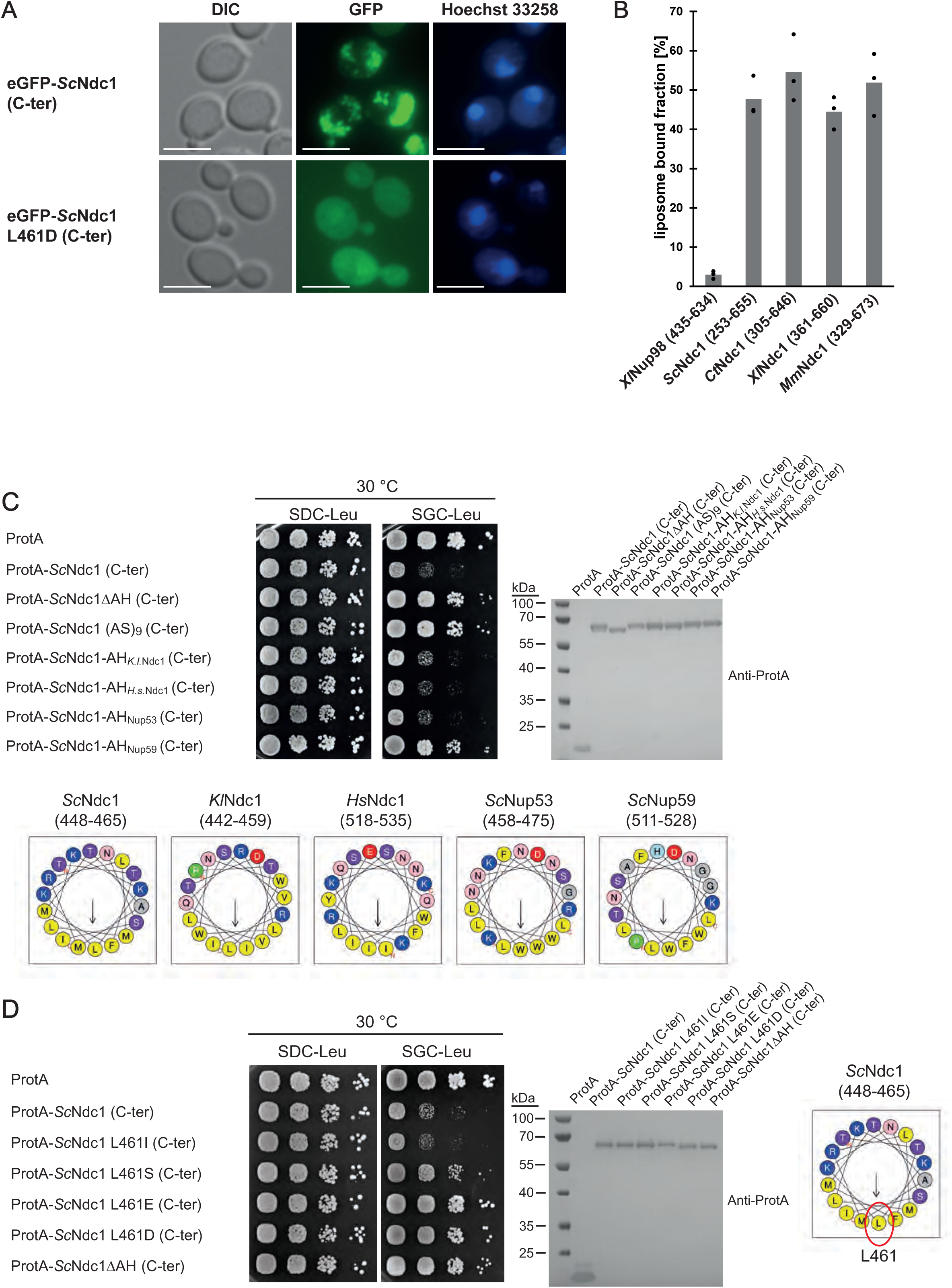
(A) Fluorescence microscopy of cells expressing eGFP-tagged *Sc*Ndc1 (261-655), (upper panels) or eGFP-*Sc*Ndc1 (261-655) additionally carrying the point mutation L461D (lower panels). Hoechst 33258 served as chromatin stain. Bars, 5 µm. (B) *In vitro* liposome binding assay using in *E. coli* expressed and purified C-termini from different Ndc1 homologs using 30 nm liposomes. Columns represent the average of three independent experiments, individual data points are indicated. A fragment of *Xenopus* Nup98 (435-634) served as negative control. (C) Growth and expression were analysed as described earlier using yeast cells expressing chimeric Ndc1 constructs comprised of the C-terminal part of *Sc*Ndc1 (ProtA-*Sc*Ndc1 (C-ter)) and different amphipathic motifs substituted for the endogenous amphipathic motif of *Sc*Ndc1 (448-465). The corresponding sequences are illustrated as helical wheel projections. As controls ProtA-*Sc*Ndc1 (C-ter) either deleted in the amphipathic stretch (ProtA-*Sc*Ndc1ΔAH (C-ter)) or instead the AH containing an 18 aa long alanine-serine spacer (ProtA-*Sc*Ndc1 (AS)_9_ (C-ter)) were expressed. (D) Growth and expression were analysed as described earlier using yeast cells expressing the C-terminus of *Sc*Ndc1 (ProtA-*Sc*Ndc1 (C-ter)) either mutated in the hydrophobic face of the amphipathic motif (as illustrated) or deleted in the complete 18 aa amphipathic stretch (ProtA-*Sc*Ndc1ΔAH (C-ter)).

**Supplemental Figure 4.**
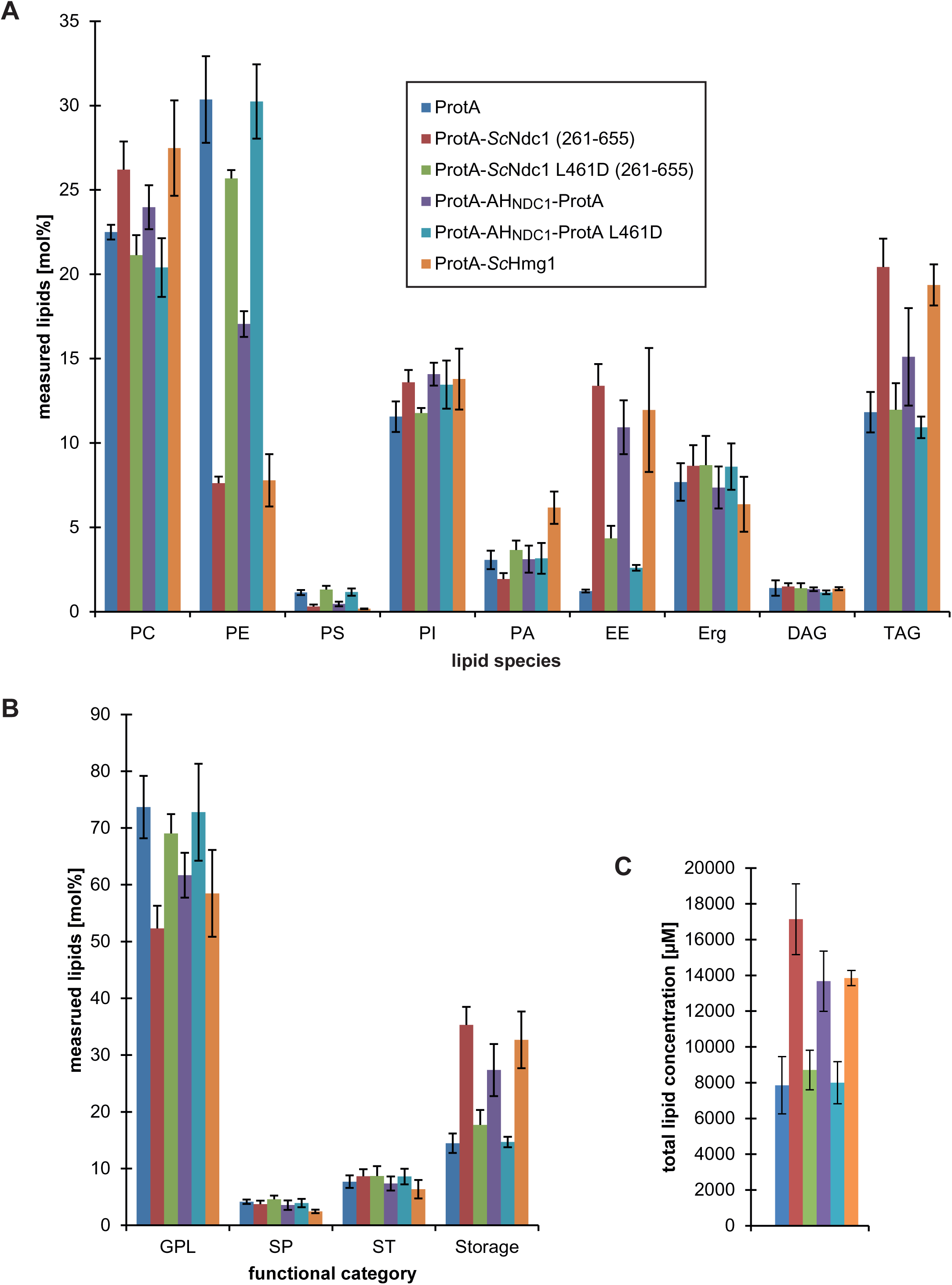
(A) Lipid profiles of cells overexpressing indicated proteins under control of the *GAL1-10* promoter. Lipids were extracted from cells grown overnight to an OD_600_ of ≈ 1.0. Lipids were subsequently analysed via nano-ESI-MS/MS. Amount of the lipid species PC (phosphatidylcholine), PE (phosphatidylethanolamine), Phosphatidylserine (PS), phosphatidylinositol (PI), phosphatidic acid (PA), ergosteryl ester (EE), ergosterol (Erg), diacylglycerol (DAG) and triacylglycerol (TAG) were plotted as mol % of measured lipids. Error bars represent standard deviation of the mean of four independent experiments. (B) Plots of the functional categories of lipids glycerophospholipids (GPL), sphingolipids (SP), sterols (ST) and storage lipids (Storage). Error bars represent standard deviation of the mean of four independent experiments. (C) Total lipid concentrations of lipid extracts prepared from equal cell numbers (OD_600_ values). Error bars represent standard deviation of the mean of four independent experiments.

**Supplemental Figure 5.**
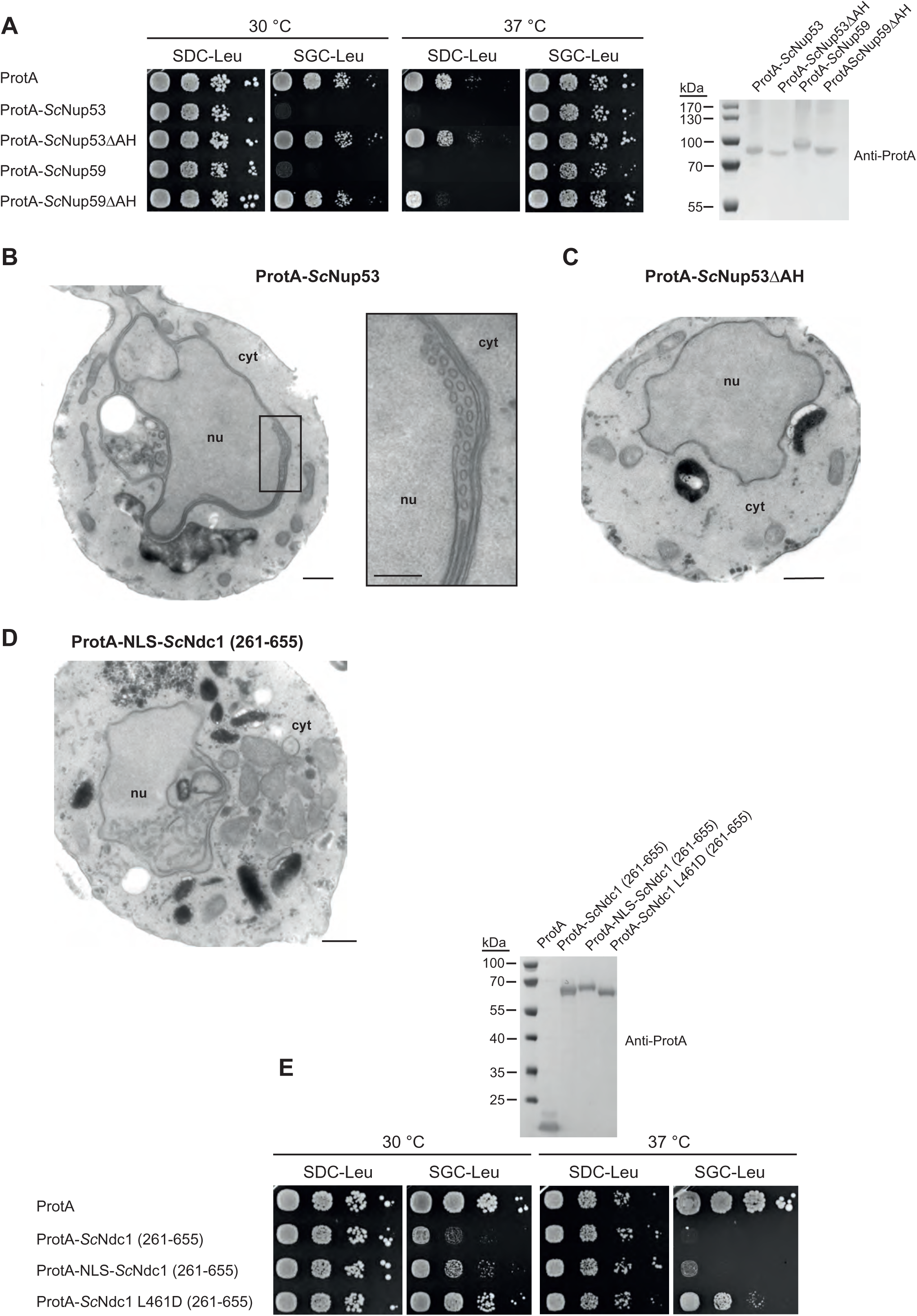
(A) Growth tests with cells overexpressing the linker nucleoporins Nup53 and Nup59 either non-mutated or mutated in their C-terminal amphipathic motifs. Growth and expression were analysed as described earlier. (B) Transmission electron microscopy of a yeast cell overexpressing ProtA-*Sc*Nup53; bar, 500 nm. Membrane sheets and tubules present at the INM are magnified; bar 250 nm. (C) Electron microscopic visualization of a yeast cell overexpressing ProtA-*Sc*Nup53ΔAH; bar, 500 nm. (D) Growth tests with cells overexpressing the C-terminus of *Sc*Ndc1 N-terminally fused to the NLS sequence of the SV40 large T-antigen. Growth and expression were analysed as described earlier. (E) Electron microscopic visualization of a yeast cell overexpressing ProtA-NLS-*Sc*Ndc1 (261-655); bar, 500 nm.

**Supplemental Figure 6.**
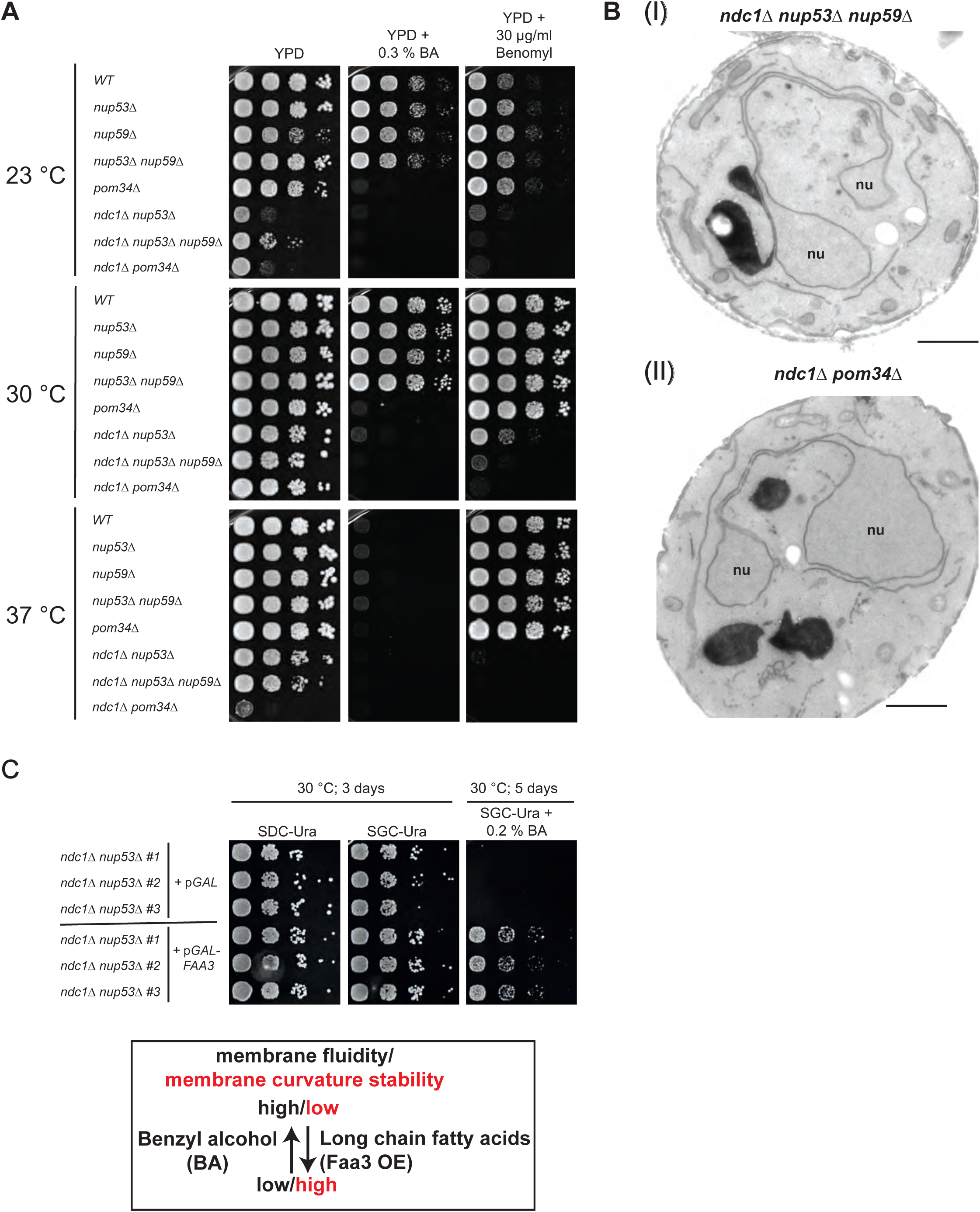
(A) Growth tests with yeast deletion strains as indicated. Growth was monitored on YPD plates using drugs either affecting membrane integrity (Benzyl alcohol) or integrity of microtubules (Benomyl) at 23 °C, 30 °C and 37 °C. Growth was analysed as described earlier. (B) Transmission electron microscopy of cells either deleted in *NDC1* and the linker nucleoporins *NUP53* and *NUP59* (a) or lacking both *Sc*Ndc1 and *Sc*Pom34 (b); bars, 1000 nm. (C) Growth tests of the *ndc1*Δ *nup53*Δ cells expressing the long chain fatty acyl-CoA synthetase *Sc*Faa3 from pYES2, an *URA3*-marker-containing 2µ plasmid, under control of the *GAL1* promoter. The *ndc1*Δ *nup53*Δ *strain* transformed with the empty pYES2 plasmid served as control. The growth tests were performed as described earlier using galactose-containing agar plates (SGC-Ura) additionally containing different amounts of the membrane-fluidizing drug benzyl alcohol. Three different transformants of each strain were used. The plates were incubated for up to 5 days at 30 °C. Possible impacts of BA and *Sc*Faa3 expression on the membrane topology and fluidity were summarized in the pictured text box.

**Supplemental Table 1.** Yeast strains used in this study

| Name | Genotype | Source/Reference |
| --- | --- | --- |
| W303-1B | *MATα ade2-1 ura3-1 his3-11,15 leu2-3,112 trp1-1 can1-100* | (Thomas and Rothstein, 1989) |
| Ndc1 shuffle | W303-1B, *ndc1::natNT2*, pRS316-*NDC1* | This study |
| Ndc1 shuffle *nup53Δ* | W303-1B, *ndc1::natNT2*, pRS316-*NDC1*, *nup53::hphNT1* | This study |
| Ndc1 shuffle nup59Δ | W303-1B, *ndc1::natNT2*,<br>pRS316-*NDC1*, *nup59::loxP* | This study |
| Ndc1 shuffle nup53Δ<br>nup59Δ | W303-1B, *ndc1::natNT2*,<br>pRS316-*NDC1*,<br>*nup53::hphNT1*, *nup59::loxP* | This study |
| Ndc1 shuffle pom34Δ | W303-1B, *ndc1::natNT2*,<br>pRS316-*NDC1*, *pom34::loxP* | This study |
| Ndc1 shuffle pom152Δ | W303-1B, *ndc1::natNT2*,<br>pRS316-*NDC1*,<br>*pom152::kanMX5* | This study |
| Ndc1 shuffle pom34Δ<br>pom152Δ | W303-1B, *ndc1::natNT2*,<br>pRS316-*NDC1*,<br>*pom34::his5<sup>+</sup>*,<br>*pom152::kanMX5* | This study |
| Ndc1 shuffle pom152Δ<br>nup53Δ | W303-1B, *ndc1::natNT2*,<br>pRS316-*NDC1*,<br>*pom152::loxP*,<br>*nup53::hphNT1* | This study |
| Ndc1 shuffle pom152Δ<br>nup53Δ pom34Δ | W303-1B, *ndc1::natNT2*,<br>pRS316-*NDC1*,<br>*pom152::loxP*,<br>*nup53::hphNT1*,<br>*pom34::his5<sup>+</sup>* | This study |
| nup53Δ | W303-1B, *nup53::hphNT1* | This study |
| nup59Δ | W303-1B, *nup59::loxP* | This study |
| nup53Δ nup59Δ | W303-1B, *nup53::hphNT1*,<br>*nup59::loxP* | This study |
| pom34Δ | W303-1B, *pom34::his5<sup>+</sup>* | This study |

**Supplemental Table 2.**
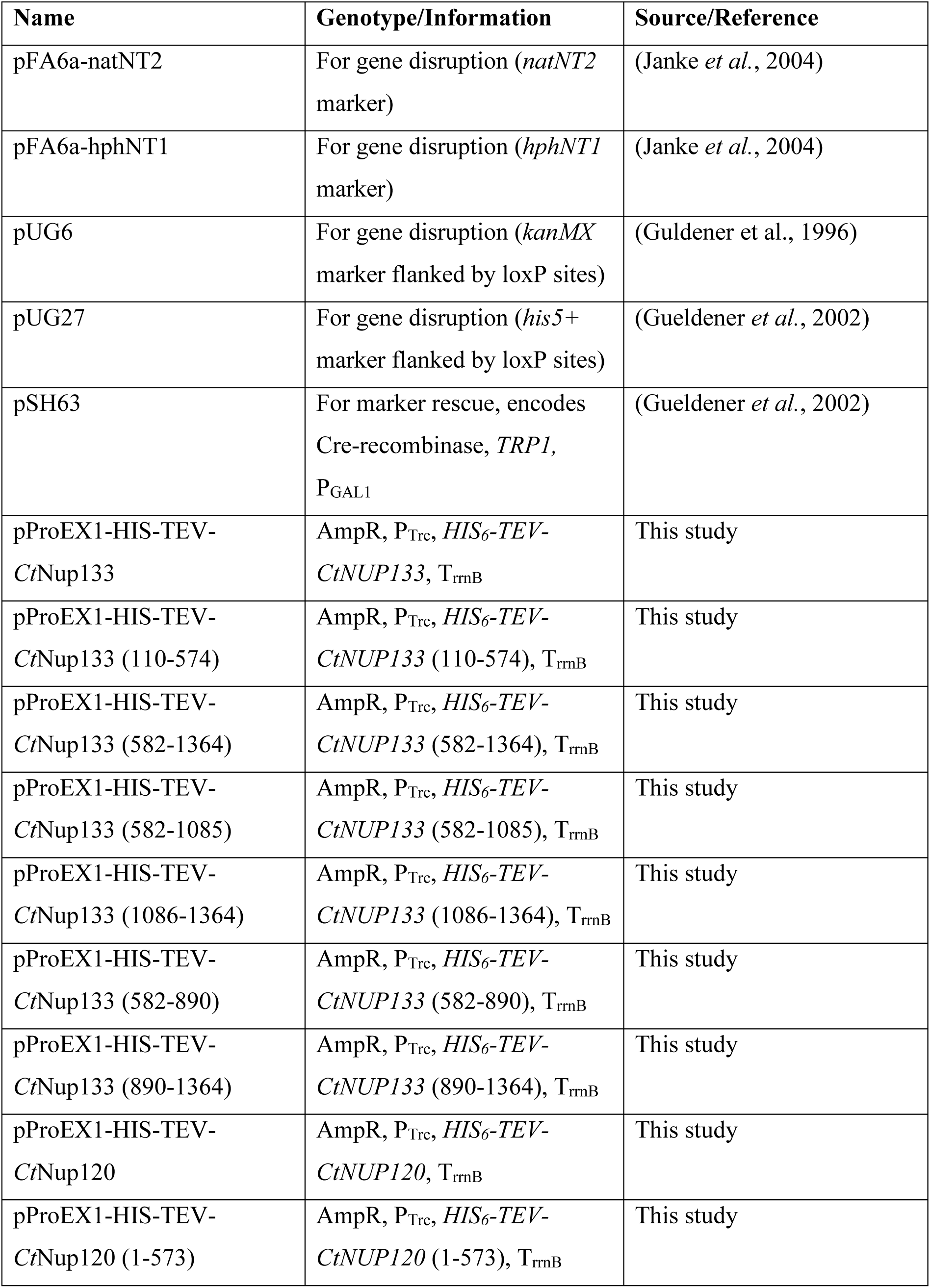
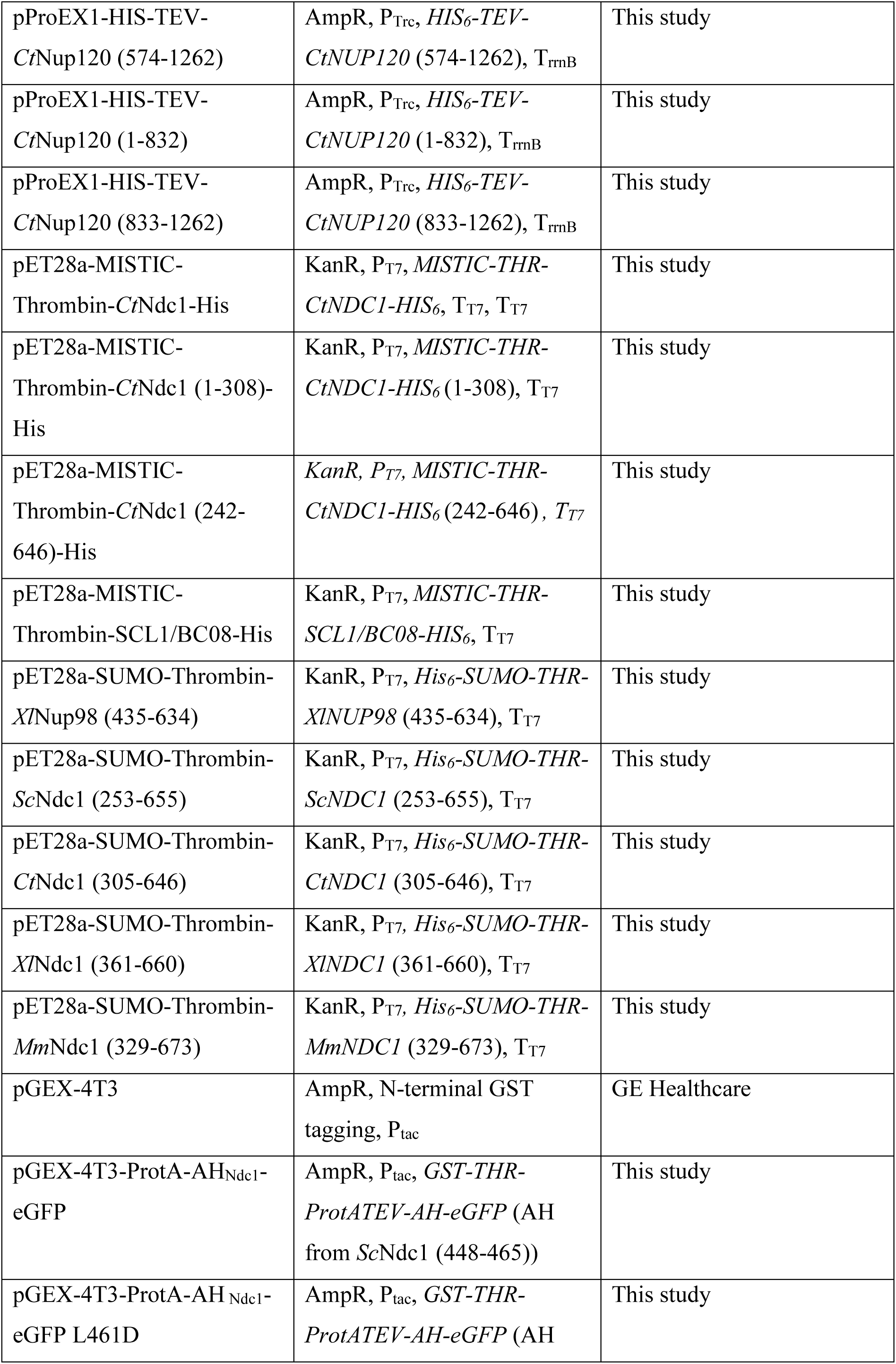
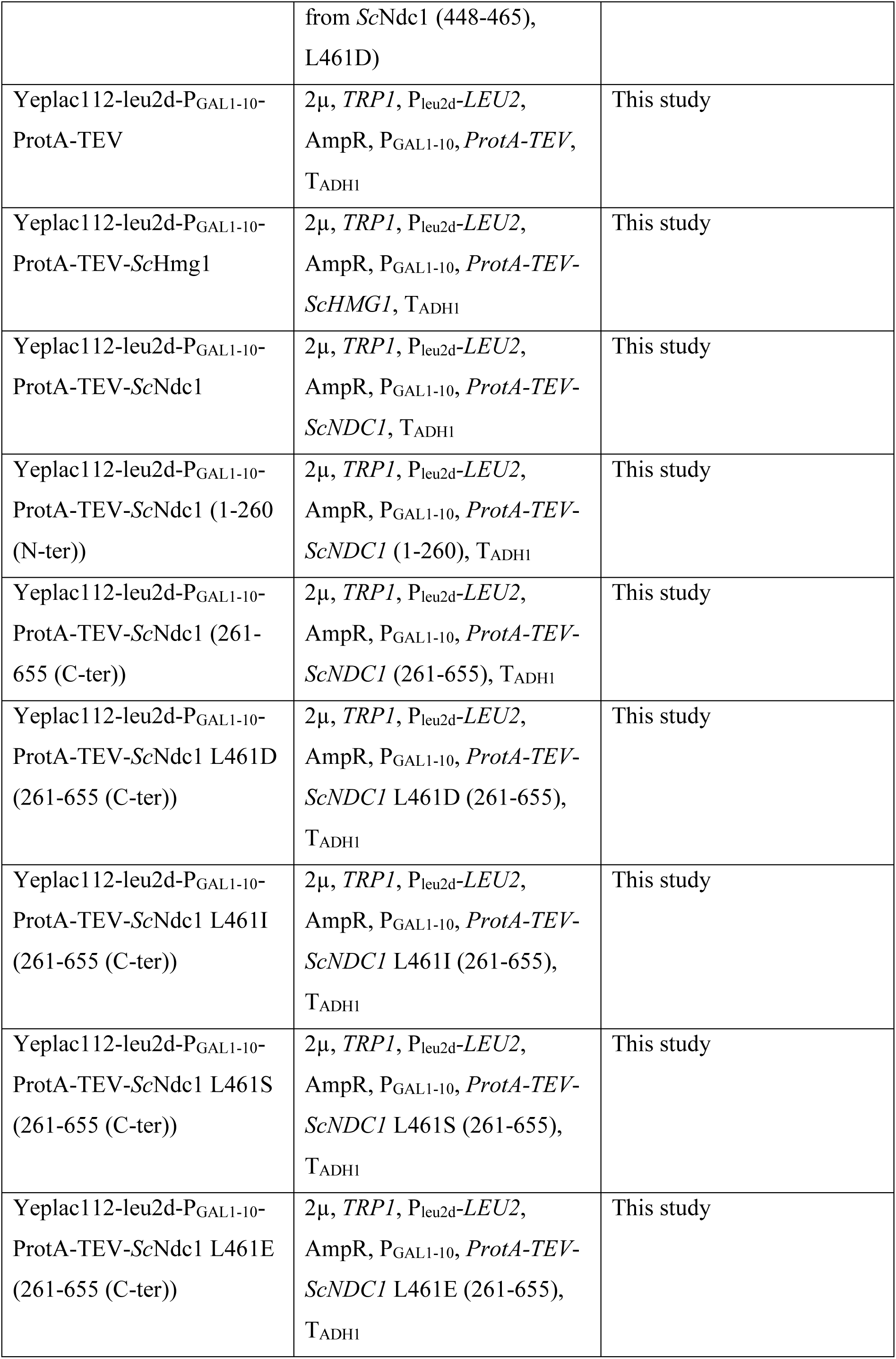
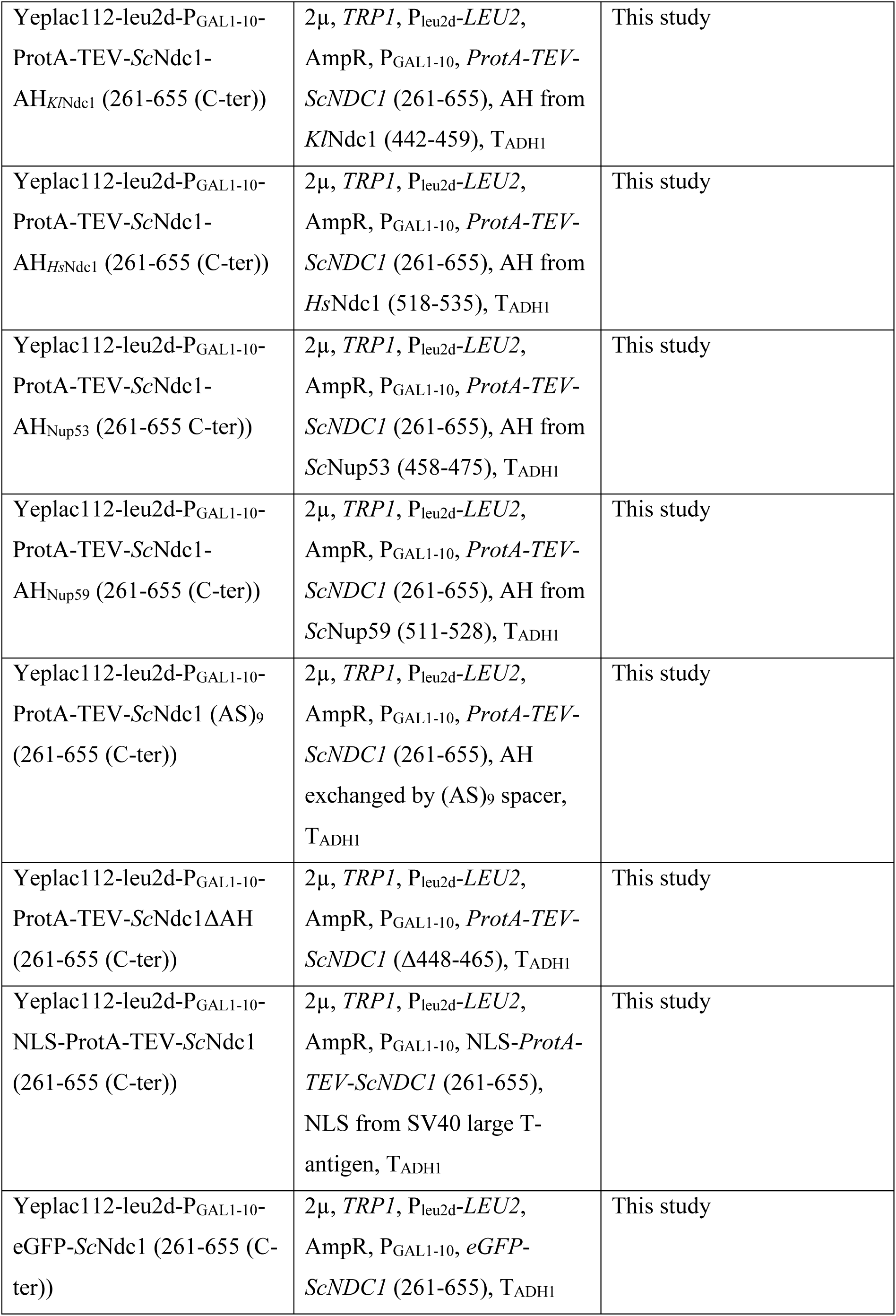
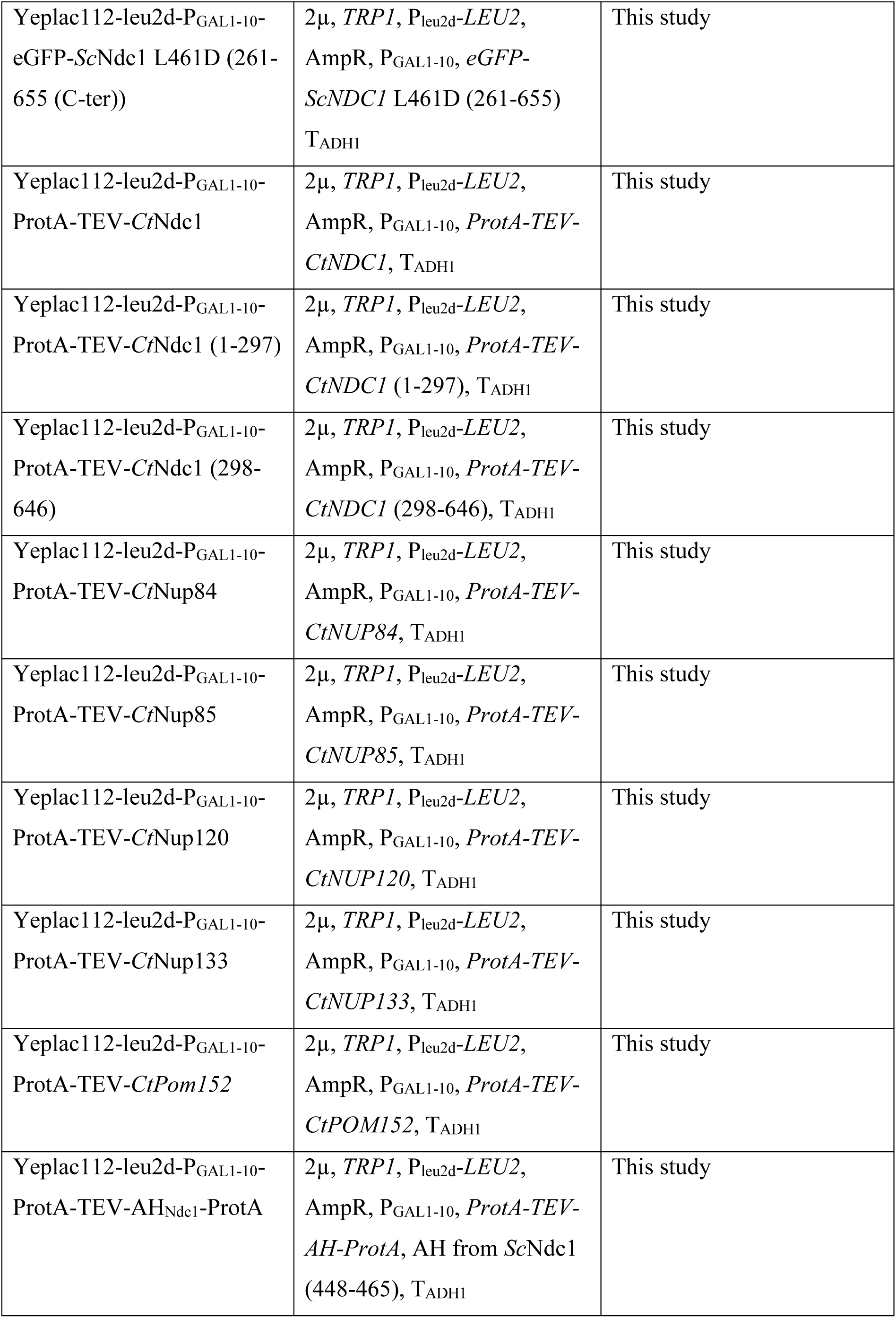
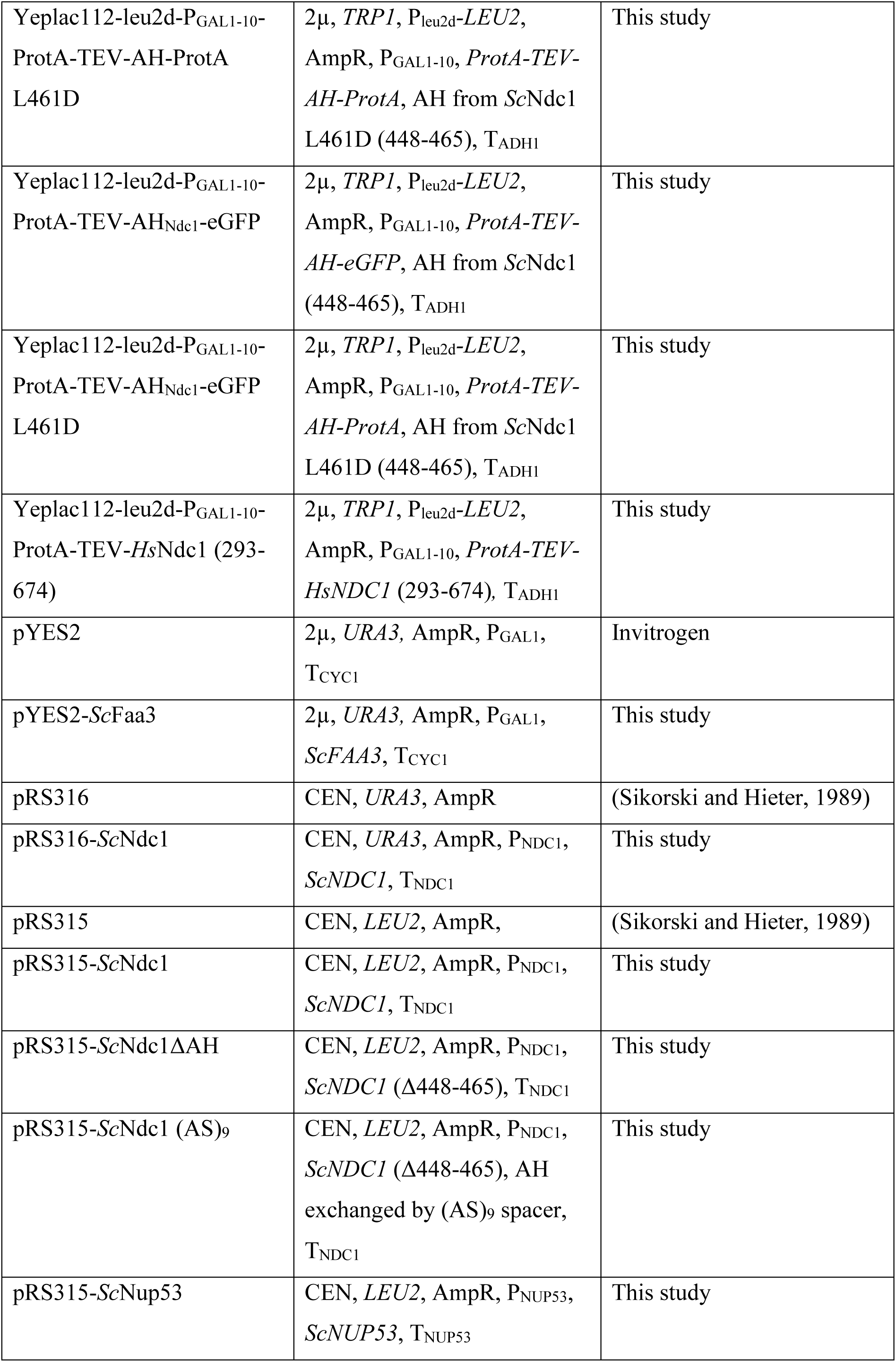
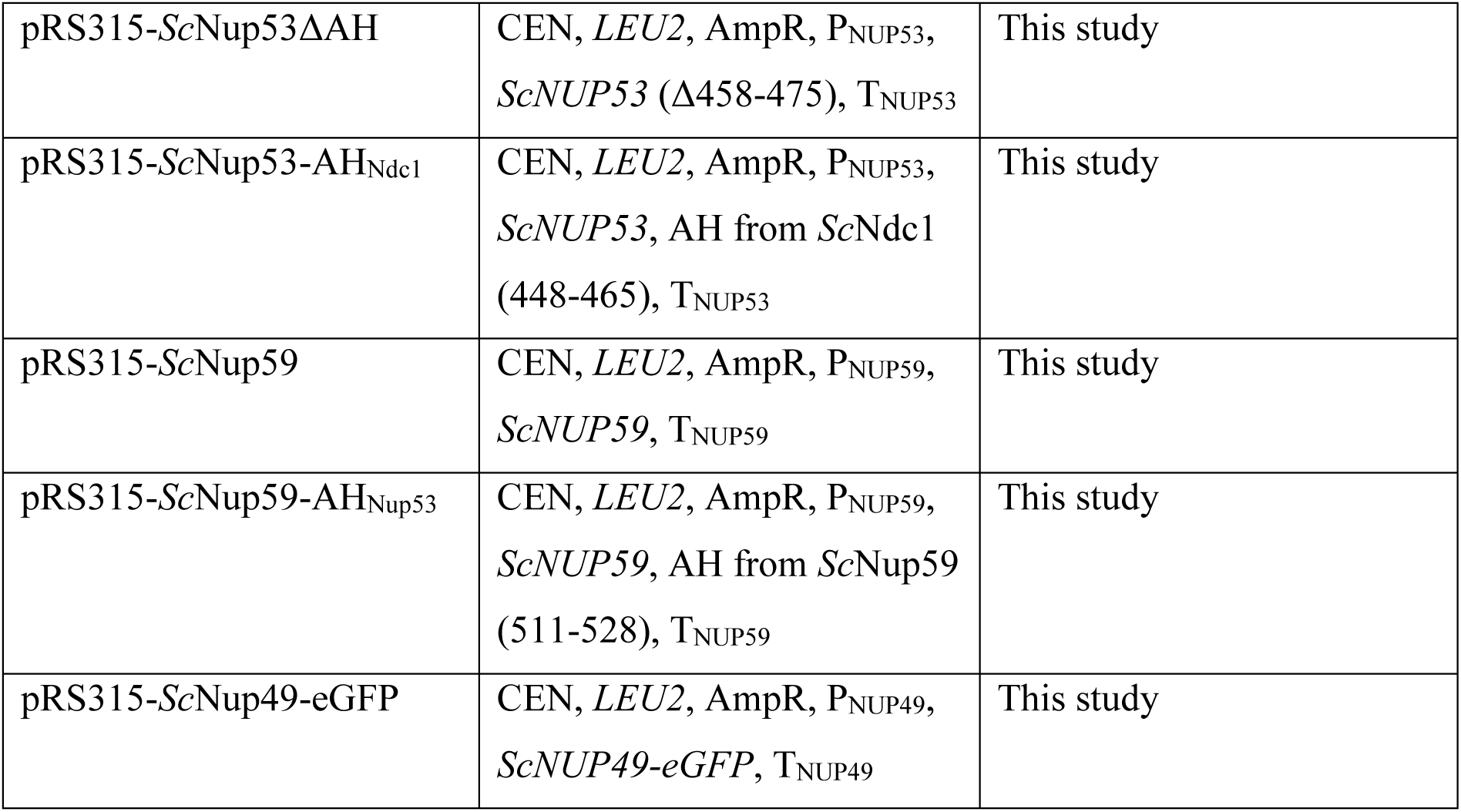
Plasmids used in this study

## AUTHOR CONTRIBUTIONS

I.A., W.A. and E.H. designed the study and analysed the data. All experiments were performed by I.A. and M.K. except TEM which was carried out by A.H., the liposome flotation assays carried out by M.T., the GUV binding assays carried out by M.W., the *in vitro* Ndc1 binding assays carried out by J.S. and the lipidomics carried out by C.L. and B.B. The manuscript was written by I.A., W.A. and E.H. All authors discussed the results and commented on the manuscript.

## ACKNOWLEDGEMENTS

E.H. and W.A. are recipients of grants from the Deutsche Forschungsgemeinschaft [Hu363/13-1, Hu363/13-2, Hu363/13-3; AN377/7-1]. We are grateful to Hilmar Bading, Department of Neurobiology and Interdisciplinary Center for Neurosciences, University of Heidelberg, for providing the opportunity to carry out the electron microscopy work in his laboratory.

## Notes

### Competing Interest Statement

The authors have declared no competing interest.

